# RUNX1-deficient human megakaryocytes demonstrate thrombopoietic and platelet half-life and functional defects: Therapeutic implications

**DOI:** 10.1101/2022.09.12.507354

**Authors:** Kiwon Lee, Hyun Sook Ahn, Brian Estevez, Mortimer Poncz

**Affiliations:** Division of Hematology, The Children’s Hospital of Philadelphia; Departments of Pathology and Laboratory Medicine and Pediatrics, Perelman School of Medicine at the University of Pennsylvania, Philadelphia, PA, 19104 USA

## Abstract

Heterozygous defects in runt-related transcription factor-1 (RUNX1) are causative of a familial platelet disorder with associated myeloid malignancy (FPDMM). Since RUNX1-deficient animal models do not mimic FPDMM’s bleeding disorder or leukemic risk, establishment of a proper model system is critical to understand the underlying mechanisms of the observed phenotype and to identify therapeutic interventions. We previously reported an *in vitro*-megakaryopoiesis system using human CD34^+^-hematopoietic stem and progenitor cells that recapitulated the FPDMM quantitative megakaryocyte defect by decreasing *RUNX1* expression using a lentiviral short-hairpin RNA (shRNA for *RUNX1* or shRX) strategy. We now show that shRX-megakaryocytes have a marked reduction in agonist responsiveness. We then infused shRX-megakaryocytes into immunocompromised NOD-SCID gamma (NSG) mice and demonstrated that these megakaryocytes released fewer platelets than megakaryocytes transfected with a non-targeting shRNA, and these platelets had a diminished half-life. The platelets were also poorly responsive to agonists, unable to correct thrombus formation in NSG mice homozygous for a R1326H mutation in von Willebrand Factor (VWF^R1326H^), which switches species-binding specificity of the VWF from mouse to human glycoprotein Ibα. A small-molecule inhibitor RepSox, which blocks the transforming-growth factor beta pathway, and which rescued defective megakaryopoiesis *in vitro*, corrected the thrombopoietic defect, platelet half-life and agonist response, and thrombus formation in NSG/VWF^R1326H^ mice. Thus, this model recapitulates the defect in FPDMM megakaryocytes and platelets, identifies previously unrecognized defects in thrombopoiesis and platelet half-life, and demonstrates, for the first time, reversal of RUNX1 deficiency’s hemostatic defects by a drug.

**Key Points:**

- RUNX1-deficient megakaryocytes exhibit thrombopoietic and platelet defects in NSG/VWF^R1326H^ mice.
- Pre-exposure of RUNX1-deficient megakaryocytes to a TGFβ1-pathway inhibitor ameliorated both defects, correcting hemostasis.

## Introduction

Lineage-specific gene regulation by key transcription factors is critical for hematopoiesis. Failures of such key transcription factors to express at the appropriate level and time can broadly affect downstream pathways of differentiation, likely at multiple levels, and the final phenotypes would involve multiple pathways in multiple lineages^1-4^. Runt-related transcription factor-1 (RUNX1) is an essential transcriptional factor in hematopoietic stem cells and is involved in multiple lineages, including megakaryocyte-formation^5-9^. *RUNX1* haploinsufficiency (RUNX1^+/-^) causes a described syndrome, familial platelet disorder with associated myeloid malignancy (FPDMM)^7,10^. Usually, patients with FPDMM have symptoms of mild-to-moderate qualitative and quantitative platelet defects with increased risk of myelodysplasia (MDS) and acute myeloblastic leukemia (AML)^11-16^.

Unlike human FPDMM, RUNX1^+/-^ mice do not have a bleeding diathesis nor do they develop either MDS or AML^6,16,17^. These studies suggests that there are significant species-specific differences in phenotypes of RUNX1 deficiency that limit usage of this non-human animal model. Without a small animal model mimicking the main features of human FPDMM, it has been a challenge to advance our understanding of functional mechanisms of the observed phenotype seen in RUNX1^+/-^ and in testing therapeutics.

As an alternative to animal models, human induced-pluripotent stem cells (iPSCs) have served as a resource for modeling human hematopoietic diseases, including transcription factors central to megakaryopoiesis^18^. RUNX1^+/-^ human iPSCs and iPSCs established from affected individuals recapitulate critical features of FPDMM patients, e.g., reducing megakaryocyte yield and functionality^19-21^. We showed that there was an accompanying depletion of megakaryocyte-biased subpopulations of hematopoietic progenitors^22^. Also, single-cell RNA sequencing data suggests that RUNX1^+/-^ induces inflammatory-related pathways such as the transforming-growth factor beta 1 (TGFβ1) signaling pathway. By blocking these pathways with a small molecule inhibitor, we observed rescue of the FPDMM phenotypes, which may have therapeutic implications^22^. Because iPSC-derived megakaryocytes are likely embryonic in nature^23^ and FPDMM is a disease affecting adult hematopoiesis, we treated CD34^+^-derived adult hematopoietic stem and progenitor cells (HSPCs) using lentiviral shRNA-targeting RUNX1 (shRX), reducing RUNX1 expression by ∼50-70%^22^. After differentiation, the resulting shRX-megakaryocytes recapitulated the decrease in megakaryocyte yield, which could be corrected by the same small molecule seen in iPSC studies.

Below, we extended these shRX studies to better understand the effects on platelet yield and on circulating platelet half-life and functionality in a murine host model. We then examined potential drug intervention to correct these defects. We began these studies *in vitro*, utilizing these shRX-megakaryocyte to examine agonist responsiveness and to examine quantitative and qualitative responsiveness of the shRX-megakaryocytes to potential therapeutics. Next, we focused on *in vivo* studies of released platelets from infused CD34^+^-derived megakaryocytes into immunocompromised NOD-scid gamma (NSG) mice, having previously shown that infused megakaryocytes release near-physiologic platelets much as endogenous^24, 25^, marrow-derived megakaryocytes release at least half of all platelets in a mouse^26^. We found that infused shRX-megakaryocytes into NSG mice released fewer platelets per megakaryocyte and the resultant platelets had a shortened half-life. Released shRX-platelets also poorly responded to agonists. To study overall hemostatic functionality of the released platelets from shRX-megakaryocytes, we used NSG mice that were homozygous for an R1326H substitution in von Willebrand factor (VWF), switching species-specificity of VWF from mice to human glycoprotein (GP) Ibα^27,28^. These mice have a bleeding diathesis unless infused with sufficient numbers of functional human platelets. We found that shRX-megakaryocyte-released platelets poorly improved hemostasis; however, shRX-megakaryocytes grown in the presence of a small-molecule inhibitor RepSox that blocks the TGFβ1 pathway^29^ improved shRX-platelet yield, half-life, and agonist responsiveness, correcting thrombus formation in a carotid artery photochemical injury model. Thus, the developed system offers a model for studies of modified human megakaryocyte and the resultant platelets, allowing detailed analysis of the released platelets, and offering a platform for preclinical screening of potential therapeutics.

## Materials and Methods

### CD34^+^ cells, ex-vivo culture and megakaryopoiesis

Mobilized human CD34^+^-derived HSPCs isolated from adult bone marrow were purchased from the Fred Hutchinson Cancer Research Center. Thawed CD34^+^ HSPCs were cultured in differentiation medium consisting of 20% BIT 9500 serum substitute (STEMCELL, 09500), 55 µM 2-mercaptoethanol (Gibco, 2198523), 40 µg/ml LDL (STEMCELL, 02698), human cytokines (1 ng/ml stem cell factor, 100 ng/ml thrombopoietin, 13.5 ng/ml interleukin 9, 10 ng/ml interleukin 6; all cytokines from R&D Systems), and 1XPenicillin-Streptomycin (Gibco, 1514022) in Iscove’s Modified Dulbecco’s Medium with GlutaMAX (Gibco, 31980030) as we previously described^22^. Cells were grown at 37°C in a 5% CO_2_ incubator for 11 to 14 days prior to study of the terminally differentiated megakaryocytes (Figure 1A).

**Figure 1.**
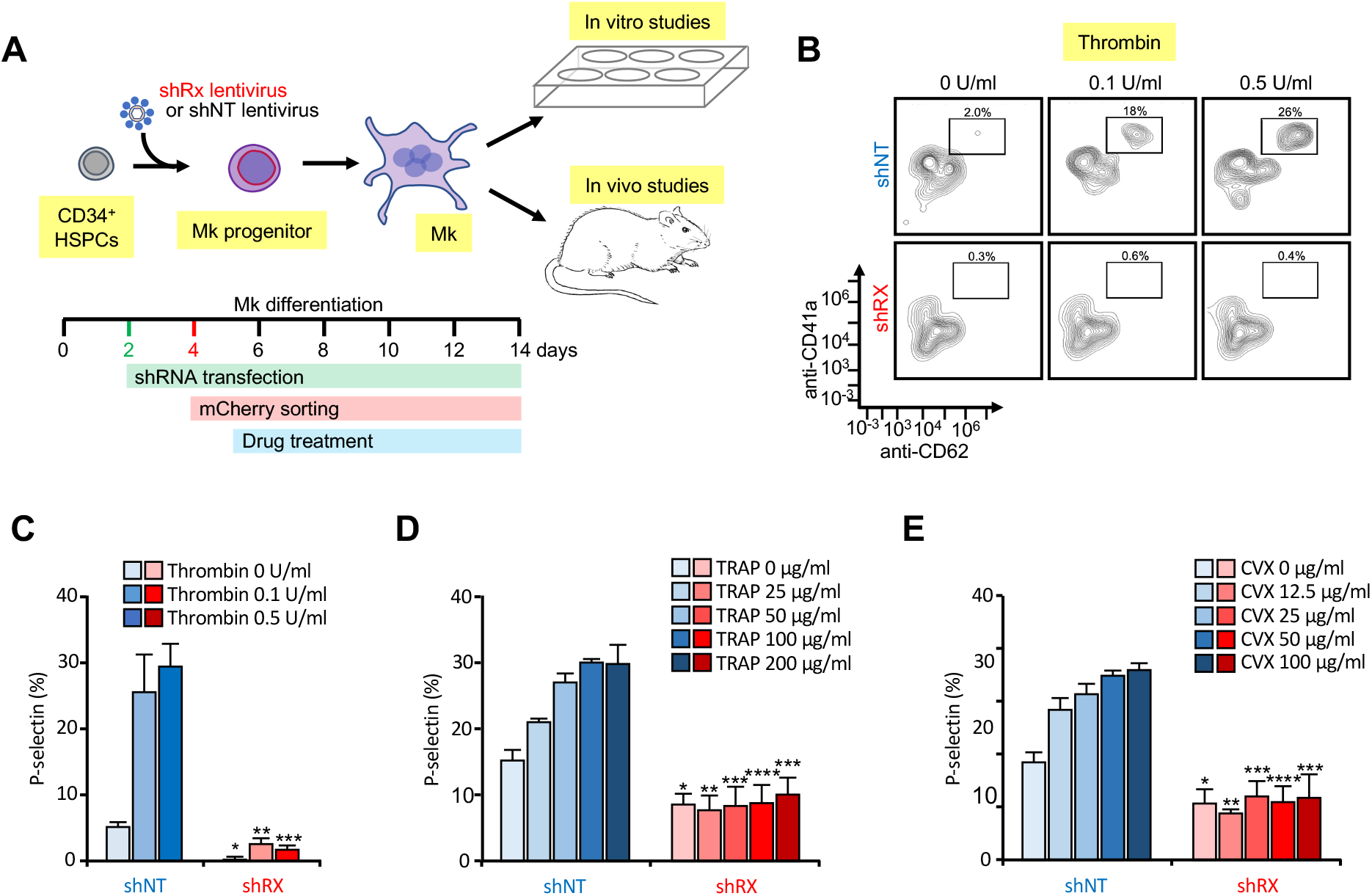
Analysis of *in vitro*-grown megakaryocytes derived from human CD34^+^ cells following RUNX1 suppression. (**A**) Experimental schema of the studies done. To mimic FPDMM disease, shRX or shNT lentiviruses were infected to CD34^+^ cells at D2 of differentiation. Infected cells expressing mCherry (mCherry^+^), sorted in Day 4 of differentiation, were the focus of these studies. From D5 of differentiation, cells were treated with drugs until Days 11 or 13-14 of differentiation. These matured megakaryocytes (Mk) were used for either *in-vitro* or *in-vivo* experiments. (**B**) Representative flow cytometric data on Day 11 of differentiation for agonist-induced surface P-selectin exposure. After stimulation of mCherry^+^ megakaryocytes with indicated doses of thrombin, cells were stained with both anti-hCD41a and hCD62P (P-selectin). (**C**) The mean ± 1 standard deviation (SD) levels of surface P-selectin were quantified in megakaryocytes stimulated by increasing doses of thrombin as indicated from lighter to heavier color. Blue = shNT-megakaryocytes. Red = shRX-megakaryocytes. In (**D**) and (**E**), similar studies as in (C), but shNT- or shRX-megakaryocytes were exposed to TRAP (D) or convulxin (E). In (C) through (E), N = 3 separate studies, each in duplicate. * = P ≤ 0.05, ** = P ≤ 0.01, *** = P ≤ 0.001, and **** = P ≤ 0.0001. P-values were calculated by one-way ANOVA comparing shRX-megakaryocyte to each shNT-megakaryocyte sample. Also, see Figure S5 for all three agonists on Day 14 megakaryocytes showing that all megakaryocytes that were mCherry^−^ for shRX lentivirus were agonist responsive.

### Lentiviral infection

At 24 hours after seeding, CD34^+^-HSPCs at 1×10^6^ per well in a 6-well plate (Ultra-low attachment surface, Corning #3471) were exposed to viral supernatant containing shRX-lentivirus (miRNA: 5’-CCTACGATCAGTCCTACCAAT-3’; see Figure Supplement (S) 1A and ^22^) or a nontargeting control (shNT) lentivirus to luciferase (miRNA: 5’-CCGCCTGAAGTCTCTGATTAA-3’; see Figure S1A and ^22^) added to the cells with 400 µg/ml of poloxamer-407 (Sigma, 9003-11-6) and incubated in a 37°C CO_2_ incubator for another 24 hours (Figure 1A). To calculate infection efficiencies, mCherry expression was measured at 72 hours post-infection using a CytoFlex 6 (Becton-Dickinson). Determination of RUNX1 mRNA levels was by qRT-PCR as described in the Supplemental Materials and Methods.

### Agonist-induced activation assays of megakaryocytes

Day (D) 13-14 differentiated CD34^+^-derived human megakaryocytes were washed in Phosphate-Buffered Saline (PBS, Gibco), and then resuspended as previously described to study agonist responsiveness^30^. A series of agonists, i.e., convulxin (Santa Cruz, SC-202554), thrombin (Sigma) or the thrombin receptor-activating peptide (TRAP, Sigma) was added to each sample and incubated at room temperature for 15 minutes. After washing the sample with PBS, allophycocyanin (APC)-labeled polyclonal rabbit anti-CD41 (BD Pharmingen, 1:100) and BV421-labeled polyclonal rabbit anti-CD62P (BD Pharmingen, 1:100) were added and activation of human megakaryocyte was determined by surface expression of P-selectin. Flow cytometric analysis was done and analyzed using FlowJo software v10.6 (Becton-Dickinson).

### Infused megakaryocyte murine studies

Immunocompromised NSG mice were used as previously described for studies of infused CD34^+^-derived human megakaryocytes to determine platelet yield and half-life^26^. At 48 hours after infection, an mCherry-positive sorted population, performed as described in the Supplemental Materials and Methods, was cultured. Equal numbers (3×10^6^/mouse) of shNT- and shRX-Day 13-14 megakaryocytes were infused via the tail vein in 200 µl of PBS over 3-5 minutes. Blood samples (∼100 µl) containing 0.38% sodium citrate were then obtained from the contralateral tail vein to measure released human platelets from the infused megakaryocytes over the subsequent 48-72 hours. Human platelets in the obtained blood were identified using APC-labeled polyclonal rabbit anti-human (h) CD41 and/or CD42b (BD Pharmingen, 1:100) and fluorescein isothiocyanate (FITC)-labeled polyclonal rabbit anti-mouse (m) CD41 (BD Pharmingen, 1:100) was used to detect murine platelets. The ratio of human to murine platelets were then determined and used to calculate yield of human platelets released per infused megakaryocytes as well as human platelet half-life. This half-life was compared to tail vein-infused donor-derived (dd) human platelets (200 × 10^6^/mouse) studied as with the infused human megakaryocytes as described^24^.

Agonist responsiveness of the released human platelets was done as described for *in vitro*-generated human megakaryocytes except on 100 µl of blood obtained by retro-orbital puncture. For *in vivo* hemostatic studies, previously generated NSG mice homozygous for VWF^R1326H^ were studied in a Rose Bengal photochemical carotid artery injury model^28^. At 4 hours after infusion of 3×10^6^ human megakaryocytes, carotid artery thrombosis was induced in mice anesthetized with sodium pentobarbital (80 mg/kg) injected intraperitoneally. Under anesthesia, Rose Bengal (75 mg/kg, Sigma) was injected via the tail veil, and the right carotid was exposed. After mounting a Doppler flow probe (Transonic Systems) on the site of carotid artery, a 540-nm laser (50 mWatts, Edmonds Optical) was used to injure the artery. Residual blood flow was monitored for an hour. After recording the blood flow data, all mice were euthanized.

### Institutional approval and statistical analysis

Studies of donor-derived platelets were from healthy donors with no bleeding/thrombotic history and not on aspirin. The Children’s Hospital of Philadelphia institutional human subjects review board approved these studies and informed consent was obtained from anonymized donors. Studies were done in accord with the Helsinki Principles. Institutional animal care and use committee approved the murine studies and animals were euthanized in accord with the American Veterinary Medical Association.

Differences between 2 groups were compared using a two-tailed Student t-test. For the multiple comparisons more than 2 groups, statistical analysis was performed by one-way Analysis of Variance (ANOVA) using GraphPad Prism version 6.07. Differences were considered significant when P < 0.05.

## Results

### Recapitulation of human FPDMM disease in in vitro-grown, CD34^+^-derived megakaryocytes

Prior studies by our group included CD34^+^-derived shRX-megakaryocytes invitro studies demonstrating a deficiency of both megakaryocyte-biased hematopoietic progenitors and terminally differentiated megakaryocytes as well as alterations in the TGFβ1-signaling pathway^22^. The overall schematic of further *in vitro* studies of these megakaryocytes studies is shown in Figure 1A. Using the described lentivirus (Figure S1) and studying sorted mCherry-positive cells, we found that mCherry positivity was preserved through differentiation with >97% of control shNT lentiviral-infected cells retaining positivity on Day 11 as did over 90% of shRX transfected cells (Figure S2). We confirmed our prior findings of a ∼50% decrease in both RUNX1 mRNA and protein levels (Figures S3A and S3B, respectively) and a >4-fold decrease in mature CD41^+^CD42^+^ megakaryocyte yield (Figure S3C). We also observed that immature CD41^−^CD42^−^ and CD41+CD42-cell populations are increased, while mature CD41^+^CD42^+^ megakaryocyte yield is markedly decreased (Figures S4A and 4B). Ploidy of mature megakaryocytes was also decreased by shRX (Figure S4C). These results are consistent with RUNX1 blocking terminal megakaryocyte differentiation. However, even with this blockage, in our differentiation system, there were few cells in other lineages (e.g., erythroid and myeloid, Figure S4B) and our focus was on megakaryocytes and released platelets.

We then extended these studies to look at megakaryocyte agonist responsiveness by flow cytometry. Agonist responsiveness studies on the megakaryocytes were done on Day 11 of differentiation of shRX-versus shNT-megakaryocytes for thrombin (Figures 1B and 1C); TRAP, which activates megakaryocytes in a PAR1-dependent manner^31,32^ (Figure 1D); and convulxin, a collagen receptor GPVI agonist^33,34^ (Figure 1E). These data are consistent with multiple receptor pathways being defective in shRX-megakaryocytes, supporting the conclusion that RUNX1 levels in CD34^+^-megakaryopoiesis are critical for not only maturation of megakaryocyte, but also for agonist-induced activation.

### Decreased human platelet yield and half-life from infused shRX-megakaryocytes in NSG mice

We next were interested in measuring *in vivo* human shRX-platelet yield per megakaryocyte, released platelet half-life and their responsiveness to agonists. We, therefore, combined the in vitro production of RUNX1-deficient megakaryocytes with our prior studies of infused human megakaryocytes releasing functional, human platelets intrapulmonary in immunocompromised mice^22,24^. We previously showed that such released human platelets in mice had similar size, half-life and agonist responsiveness to that of freshly drawn, healthy dd-platelets^24^. We measured both quantity and quality of human platelets that were released from infused shRX-megakaryocytes compared to shNT-megakaryocytes and dd-platelets. We infused 3×10^6^ of uninfected or shNT- or shRX-lentiviral infected megakaryocytes or 4×10^8^ dd-platelets into NSG mice. We monitored circulating human platelets by flow cytometry for 24 hours-post infusion using hCD41^+^:mCD41^+^ ratio as a measure of circulating human platelets in the recipient mice (Figure 2A). Infused dd-platelet numbers peaked almost immediately. After infusion of uninfected megakaryocytes, circulating human platelets peaked over the next 4-6 hours as we had described^24^ This delay is likely due to the time needed to release platelets from megakaryocytes entrapped in the lungs^24,26^. We next infused the same number of mCherry^+^ shNT- and shRX-megakaryocytes into NSG mice (Figure 2B). The release of platelets from shNT-megakaryocytes was similar to uninfected megakaryocytes with a peak platelet count at 4-6 hours after megakaryocyte infusion. In contrast, the number of circulating human platelets in the mice infused with a similar number of shRX-megakaryocytes was ∼30% of the shNT-platelet number and peaked earlier at ∼2 hours.

**Figure 2.**
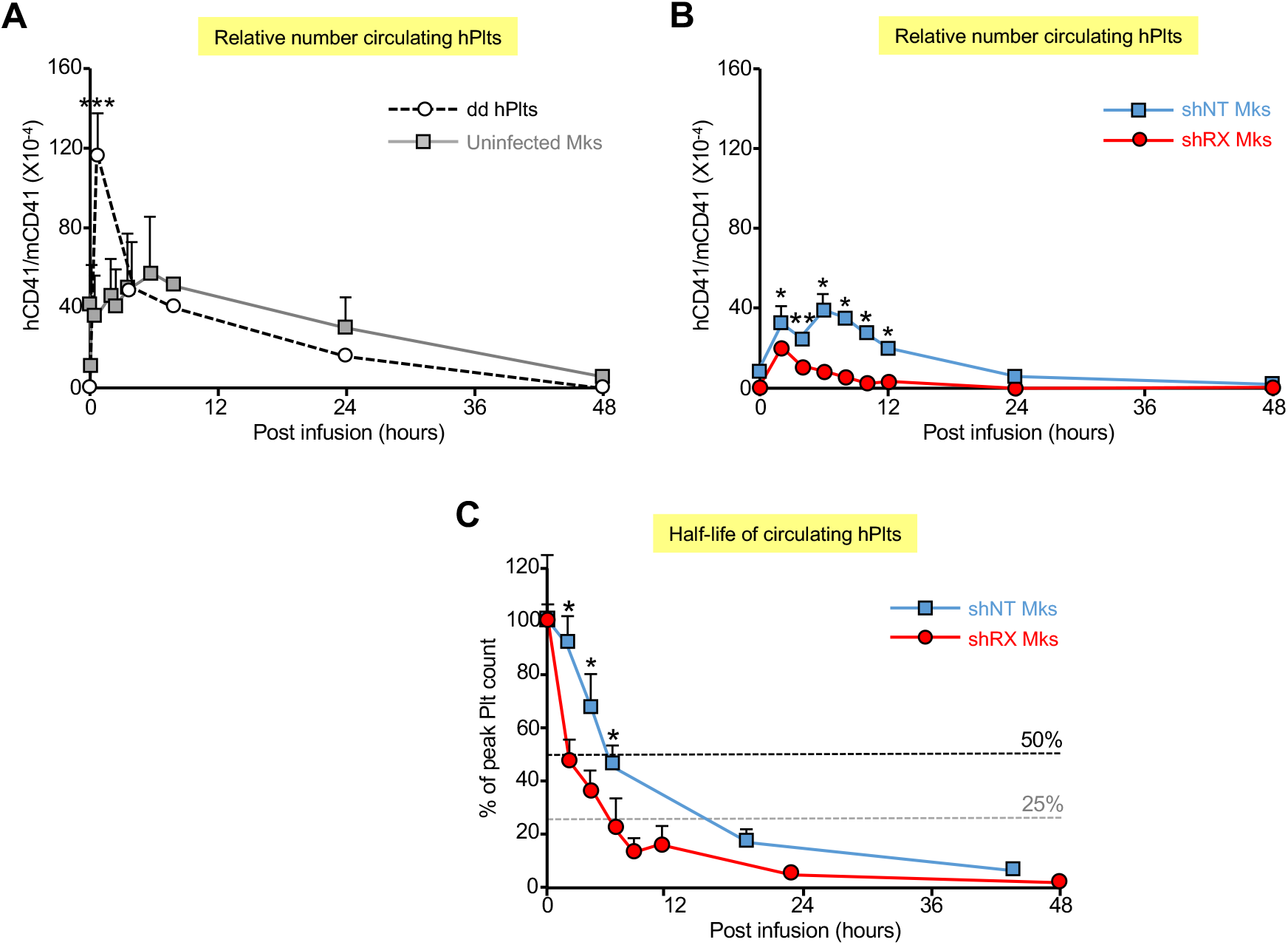
Number of circulating human platelets and their half-life released from infused dd-platelets and megakaryocytes into NSG mice. (**A**) 4×10^8^ human dd platelets or 3×10^6^ CD34^+^-megakaryocytes were infused into NSG mice. At each time point, peripheral blood was withdrawn to measure circulating human platelet number (hPlts) relative to murine platelets after staining with hCD41 and mCD41 antibodies. Mean ± 1 SD is shown. N = 3 per arm. *** = P ≤ 0.001 by one-way ANOVA compared to CD34^+^-megakaryocytes samples. (**B**) Same as in (A), but for 9 X10^6^ infused mCherry^+^ shNT- and shRX-megakaryocytes. N = 3 per arm. * = P ≤ 0.05 and ** = ≤ 0.01 by one-way ANOVA comparing shNT-platelets released vs. shRX-platelets. (**C**) Similar to (B), but after 9×10^6^ of shNT- or shRX-megakaryocytes were infused and shown to show drop from peak platelet count with the 50% and 25% levels indicated. N = 3 per arm. * = P ≤ 0.05 by one-way ANOVA comparing shNT-platelets released vs. shRX-platelets.

To compare platelet half-life of shNT- and shRX-platelets, we increased the number of infused shNT- and shRX-megakaryocytes to 9×10^6^ per recipient mice and monitored the mice more frequently and for up to 48 hours (Figure 2C). An initial half-life for shRX-platelets was ∼2 hours compared to shNT-platelets of ∼6 hours in the first 6 hrs after reaching peak platelet count (p < 0.01), and a subsequent half-life for shRX-platelets was ∼5 hours compared to ∼12 hours for shNT-platelets. The shorter initial half-life of shRX-platelets likely impacted the time and the height of the peak of released platelets relative to shNT-platelets. The basis for the shortened half-life is unclear, and whether it involves intravascular or extravascular destruction in the liver or spleen of the human platelets is also unclear at present.

### Decreased agonist responsiveness by released shRX-platelets in NSG mice

To examine whether circulating human shRX-platelets are responsive to agonist, we isolated total platelets from the peripheral blood of NSG mice infused with dd-platelet or uninfected, shNT- or shRX-megakaryocytes 2-hours post-infusion. TRAP, which is human platelet-specific^35^, was added to the samples at increasing concentrations. In line with our observation of *in vitro* megakaryocyte activation with agonists shown in Figure 1D and Figure S5C, circulating shRX-platelets had impaired activation in response to TRAP (Figure 3A).

**Figure 3.**
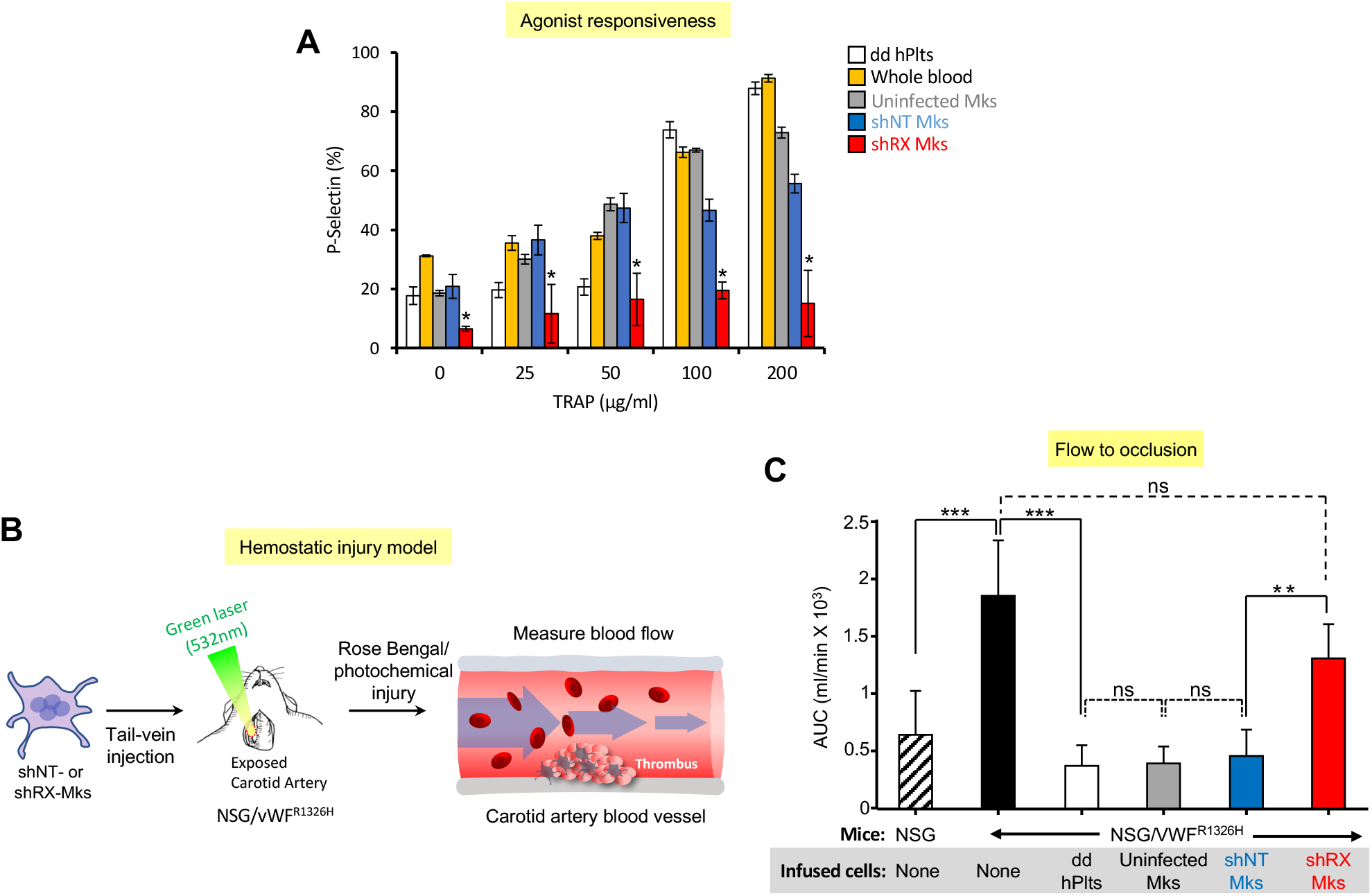
Studies of released human platelets: agonist responsiveness and hemostatic efficacy. (**A**) Flow cytometric studies of removed murine blood at two hours after infusion with dd-platelets or uninfected, shNT-, or shRX-megakaryocytes into NSG mice for P-selectin levels following activation with various concentrations of TRAP. Mean ± 1 SD is shown. N = 3 per arm. * = P ≤ 0.05 by one-way ANOVA comparing shNT-platelets released vs. shRX-platelets. (**B**) Schematic of the Rose-Bengal photochemical carotid artery thrombotic challenge with NSG/VWF^R1326H^ mice and infused human megakaryocytes or platelets. (**C**) Same as in (A) except that NSG mice were studied as a hemostatic control as the untreated NSG/VWF^R1326H^ mice had a hemostatic defect. Studies were done four hours after infusion of human platelets or megakaryocytes. Mean ± 1 is shown for residual blood flow following carotid artery injury. N = 4-6 animals per arm. ** = P ≤ 0.01, *** = P ≤ 0.001, and ns = not significant by one-way ANOVA.

### Impaired thrombus formation by infused shRX-megakaryocytes in NSG/VWF^R1326H^ mice

To better study the hemostatic effects of the various quantitative and qualitative defects in RUNX1-deficient megakaryocytes and platelets, mice were infused into NSG/VWF^R1326H^ mice with dd-platelets or uninfected or shNT- or shRX-megakaryocytes and monitored during thrombus formation for residual blood flow following a Rose-Bengal-photochemical carotid artery injury (Figure 3B). We generated these NSG/VWF^R1326H^ mice previously to characterize human platelet function *in vivo*^28^. These NSG mice are homozygous for VWF^R1326H^, leading to a switch in species specificity of binding by the VWF from mouse to human platelets. NSG/VWF^R1326H^ mice have a bleeding diathesis compared to NSG mice if not infused with functional human platelets (Figures 3C). In the various arms of this study, initial blood flow was nearly identical (Figure S6), supporting the concept that animals were studied under similar conditions. Infusion of 3×10^6^ uninfected or shNT-megakaryocytes into NSG/VWF^R1326H^ mice was sufficient to decrease the time to occlusion and total blood flow to that seen in NSG mice and in NSG/VWF^R1326H^ mice after infusion of 3×10^8^ dd-platelets or 3×10^6^ shNT-megakaryocytes (Figure 3C). In contrast, infusion of 3×10^6^ shRX-megakaryocytes had little decrease in total blood flow.

### Drug screening of shRX-HSPCs to correct megakaryocyte yield and agonist responsiveness

We screened several lead drugs and pathways that may improve outcome in RUNX1^+/-^ both for megakaryocyte yield and agonist responsiveness^21,22,36^. On D5 of differentiation (Figure 1A), shNT- or shRX-infected cells were exposed to each drug at doses based on prior studies^21,22,29,37-46^. We focused our studies of yield and function on Days 11 and 14 megakaryocytes (Figure S7 and 4, respectively) as these days of differentiation megakaryocytes have both been studied in other drug-screening studies^22,30,47,48^. RUNX1 deficient-megakaryocytes, without drug treatment, resulted in a decrease of megakaryocyte yield to ∼20% compared to shNT controls at both Days 11 and 14. At both timepoints, most tested drugs showed either modest enhancements of megakaryocyte yield or no effect on shNT-megakaryocytes yield, while treatment of shNT-HSPCs with 1 and 10 µM Ruxolitinib suppressed megakaryocyte yield (Figures 4A and S7A). In studies of shRX-megakaryocyte, the TGFβ pathway inhibitor RepSox was the only compound that improved outcome up to that seen in control cells for both days of differentiation (Figures 4B and S7B). The NOTCH pathway inhibitor DAPT and Avagacestat, JNK2 inhibitors JIN8 and JIN10, and the mTOR inhibitor MHY had more modest positive effects on Day 14, not apparent on Day 11. RepSox reversed the blockade in megakaryocyte differentiation with a decrease in observed CD41^−^CD42^−^ and CD41^+^CD42^−^ cells on Day 11 of differentiation as well as increasing the number of mature megakaryocytes and returning ploidy to that seen in shNT-megakaryocytes (Figures S3C and S4).

**Figure 4.**
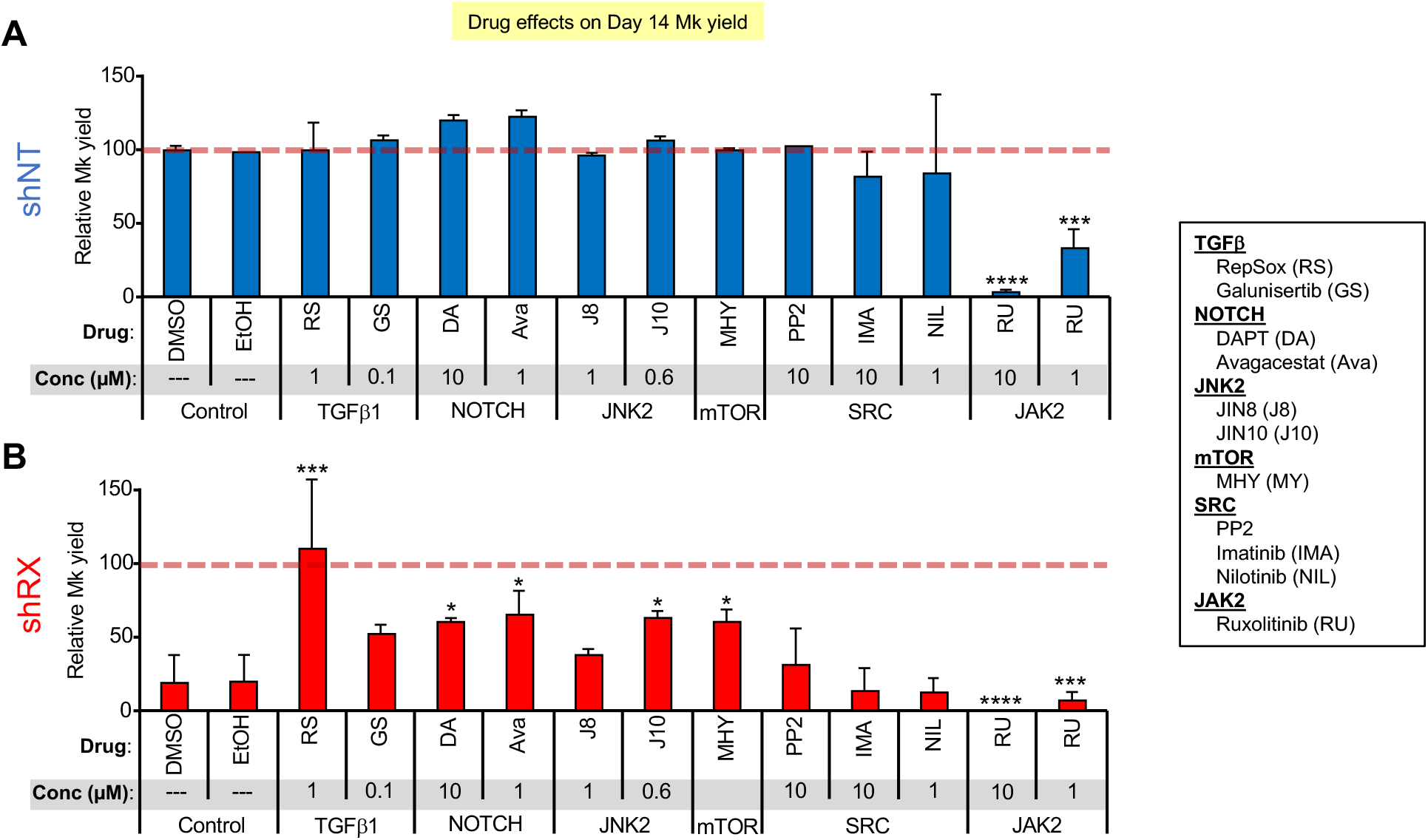
Drug screening to rescue RUNX1-deficient megakaryocyte yield. (**A**) and (**B**) are studies of shNT- and shRX-megakaryocytes, respectively, exposed to the indicated drugs from Day 5 until 14. Megakaryocyte yield was calculated by flow cytometric analysis stained for an anti-hCD42b antibody and for mCherry. The dashed line represents yield of megakaryocytes from uninfected HSPCs not exposed to any drug. Mean ± SD are shown. N = 3-5 separate studies each in duplicate. * = P ≤ 0.05, *** = P ≤ 0.001, and **** = P ≤ 0.0001 by one-way ANOVA compared to each DMSO control sample.

We then tested small drugs (RepSox, Galunisertib, DAPT, JIN10, and Avagacestat) that had moderate or greater effects in our assay systems or in prior studies^21,22,38,44,46^ and examined agonist-induced megakaryocyte responsiveness to various concentrations of TRAP by measuring P-selectin expression on Day 11 (Figure 5). Studies of agonist responsiveness in megakaryocytes have been done by many groups since first described in an analysis of integrin intracellular signaling compared to platelets^49^. Day 11 shNT-megakaryocytes had a high baseline level of P-selectin expression that further increased with exposure to TRAP (Figure 5A), while shRX-megakaryocytes had a lower baseline P-selectin expression level, and a minimal increase after TRAP (Figure 5B). Of the various drug exposures, RepSox treatment had the greatest effect on TRAP response, increasing it near to that seen with shNT-megakaryocyte (Figure 5A vs. 5B).

**Figure 5.**
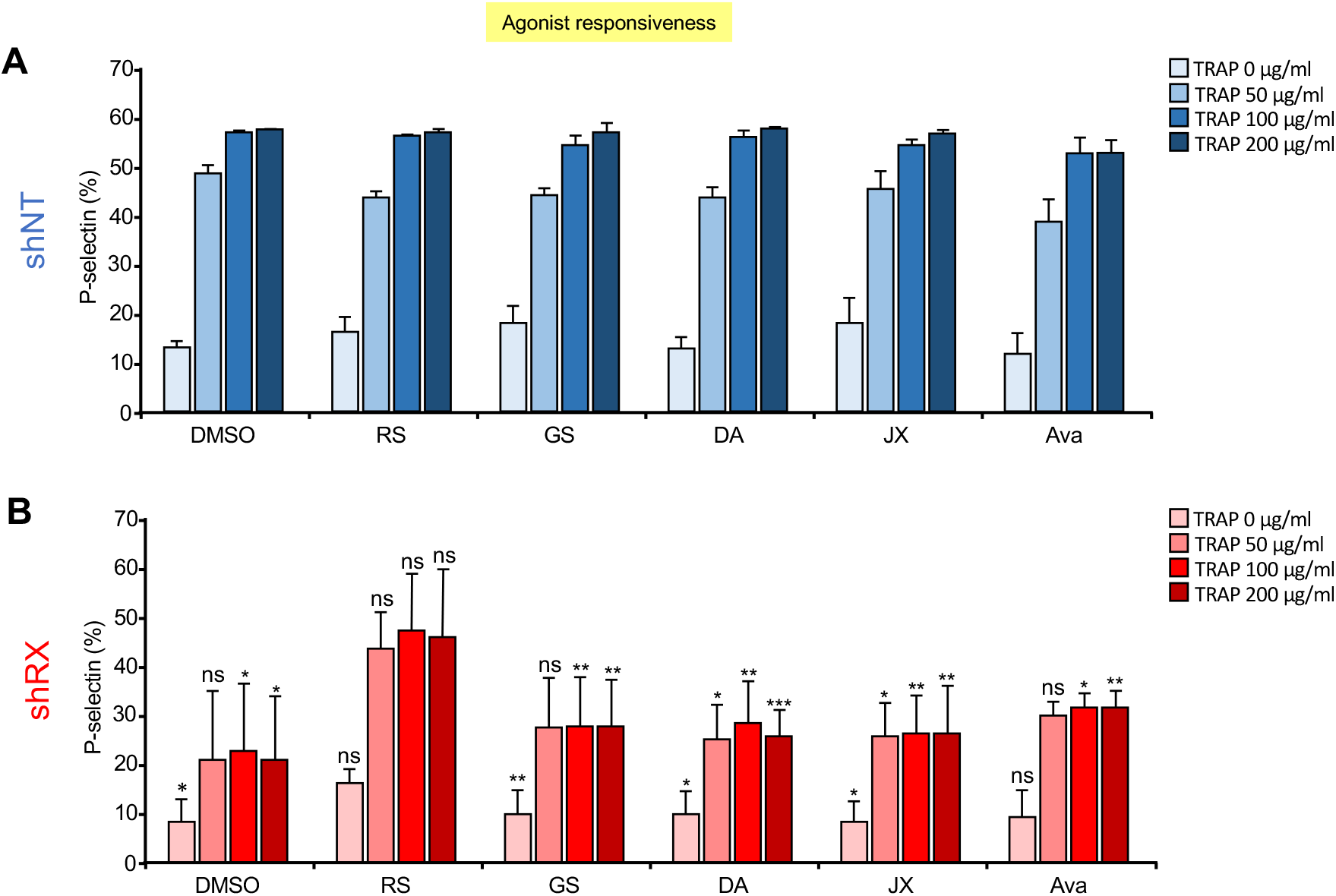
Drug screening to rescue RUNX1-deficient megakaryocyte agonist response. shNT-(**A**) and shRX-megakaryocytes (**B**) were treated with the indicated drugs from Day 5 until 11 of differentiation. Day 11 megakaryocytes exposed to various concentrations of TRAP as indicated and measured the expression of P-selectin levels by flow cytometry after staining with human CD62P. Mean ± 1 SD is shown. N = 3. * = P ≤ 0.05, ** = P ≤ 0.01 and *** = P ≤ 0.001 by one-way ANOVA comparing each shRX data point to its shNT comparative.

### RepSox as a model of therapeutic intervention in RUNX1^+/-^

We propose a model using shRNA suppression of RUNX1 in CD34^+^-HSPCs with subsequent differentiation into megakaryocytes followed by their infusion into NSG/VWF^R1326H^ mice as a model that would facilitate testing of potential therapies to correct the platelet-related bleeding diathesis in patients with FPDMM (Figure 3B). Of the various drugs tested, RepSox was most effective at correcting shRX-megakaryocyte yield and agonist responsiveness (Figures 4, 5, and S7). When shRX-HSPCs were differentiated in the presence of RepSox and then infused into mice, the drug partially to completely corrected platelet yield and half-life, and agonist responsiveness (Figures 6A and 6B, respectively). We then asked whether RepSox-exposed shRX-megakaryocytes infused into NSG/VWF^R1326H^ mice would correct the bleeding diathesis in the photochemical carotid artery injury model and observed near-complete correction of both total blood flow until occlusion and time to occlusion (Figures 7A and 7B, respectively).

**Figure 6.**
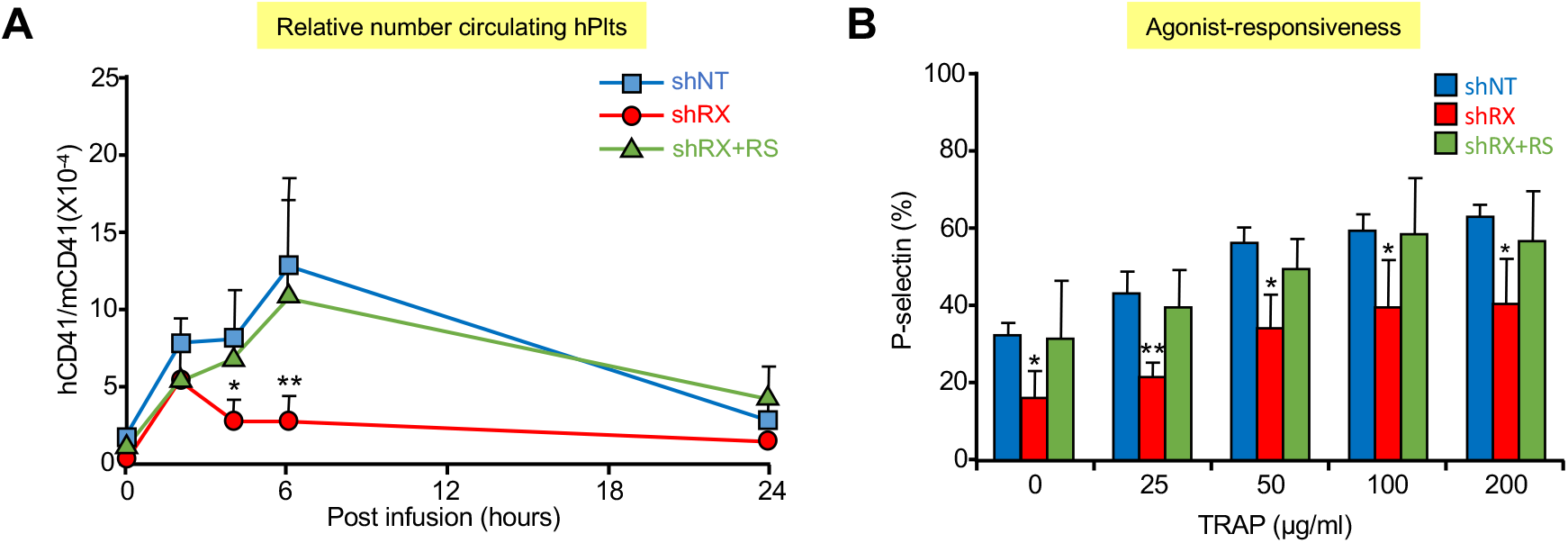
Drug correction effects of defects on platelet release and function with RepSox. (**A**) 3×10^6^ shNT- or shRX- or shRX+RepSox (shRX+RS)-megakaryocytes were infused into NSG mice. At each time point, mouse peripheral blood was withdrawn to monitor human platelet level. The peripheral blood samples stained with hCD41 and mCD41 antibodies and analyzed by flow cytometry for human vs. mouse platelets as in Figure 2. Mean ± 1 SD are shown. N = 3 per arm. * = P < 0.05 and ** = P < 0.001 comparing shRX vs. shRX+RS studies. (**B**) Study as in (A). P-selectin levels on released human platelets in mouse blood were measured under activation with various concentration of TRAP by flow cytometry. N = 3 per arm. * = P ≤ 0.05 and ** = P ≤ 0.001 by one-way ANOVA comparing shRX vs. shRX+RS studies.

**Figure 7.**
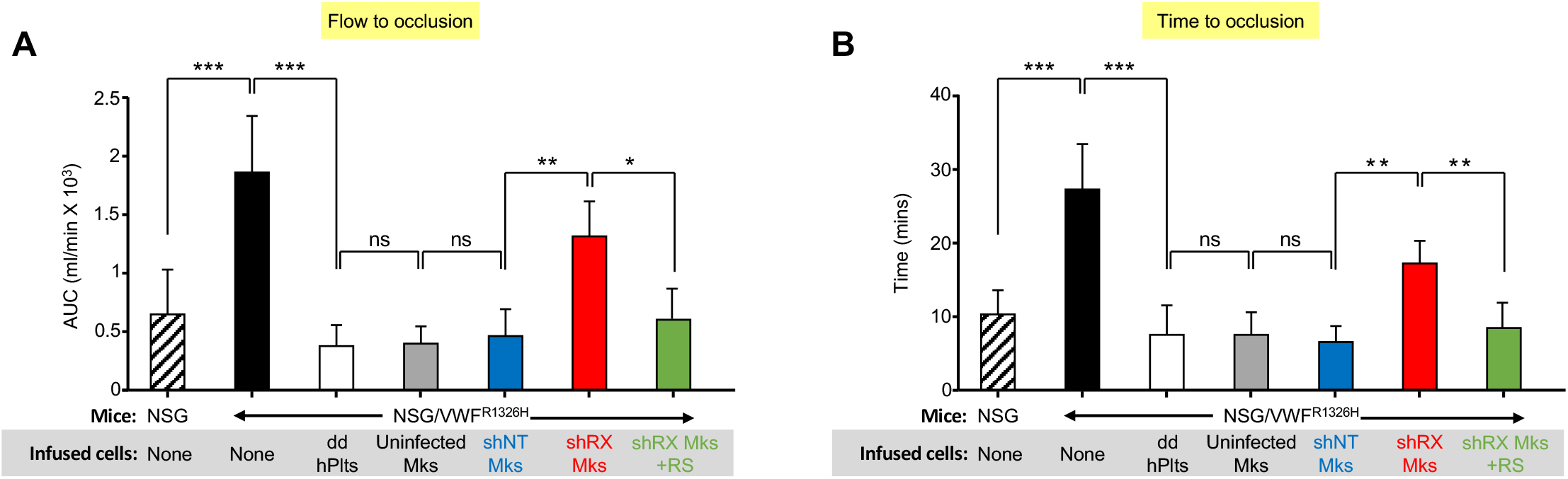
*In vivo* hemostatic correction of shRX-platelets by RepSox exposure of developing megakaryocytes. Thrombus formation studies as in Figure 3 with infused shRX-megakaryocytes in a Rose-Bengal-photochemical carotid injury model system in NSG/VWF^R1326H^ with NSG mice used as a positive control. To determine in vivo functionalities of infused RepSox-treated, shRX-megakaryocytes, we monitored thrombus formations by measuring total blood flow (**A**) and time to occlusion (**B**). Mean ± SD are shown. N = 4-6 per arm. * = P ≤ 0.05, ** = P ≤ 0.01, *** = P ≤ 0.001, and ns = not significant by one-way ANOVA comparing indicated matches.

## Discussion

The bleeding manifestations in FPDMM are often present since birth and lead to life-long risk of acute bleedings, chronic injuries like joint deformities, and even death^7,10,50^. Moreover, while patients’ platelet counts are often >50,000/µ^3^, patients often require platelet transfusions to prevent traumatic and surgical bleeding. There are challenges associated with treating patients with FPDMM: their higher baseline platelet count interferes with the physician’s ability to estimate response to a platelet transfusion as opposed to the more common cause of acquired platelet disorders associated with severe thrombocytopenia. This paper clearly delineates that RUNX1 deficiency is associated with a combination of significant qualitative and quantitative defects in the megakaryocytes and released platelets that affects multiple pathways and result in pleiotropic defects. Thus, while the focus of finding therapeutic interventions for patients with FPDMM has been on its leukemic propensity, which approaches 50% by the age of 40 years^51^, the bleeding manifestation is a significant medical challenge in these patients as well and a better understanding of the pathogenesis of these defects and therapeutic response is needed.

Our studies provide insights into the defects in the megakaryocyte:platelet axis in individuals with RUNX1 deficiency. Not only is there the well-recognized defect in megakaryocyte yield, but in the ability of these megakaryocytes to release platelets from entrapped megakaryocytes in the pulmonary bed in recipient immunocompromised mice. The released platelets have a decreased half-life as well. Thus, there are at least three reasons for the observed thrombocytopenia in patients with FPDMM: 1) defects in megakaryopoiesis, 2) thrombopoiesis, and 3) platelet half-life. Neither the importance of each defect in the final thrombocytopenia nor the mechanistic bases of each defect has yet to be defined. Moreover, RUNX1 haploinsufficiency also decreases responsiveness to multiple agonists. The finding that multiple agonist receptor pathways are defective in RUNX1-deficient megakaryocytes:platelets is not surprising given the key role this transcription factor has in both hematopoiesis and megakaryopoiesis^5,6^. It would also not be surprising that the observed phenotype varies not only based on the degree of residual effective RUNX1, but likely also by different susceptibility of various pathway to low RUNX1 levels based on an individual’s genotypic background.

Unlike developing therapeutics intended to decrease the long-term risk of leukemic progression, drug correction of the qualitative and quantitative platelet defects in FPDMM should have rapid onset of action. Measurements of platelet counts and platelet aggregation are standard clinical laboratory studies, and a short therapeutic intervention should ideally generate platelets that circulate and maintain function for days-to-weeks, be fully functional and thus be useful for hemostatic prophylaxis. We envision short-term peri-operative drug treatment may be a useful first application of such platelet-correcting drugs in patients with FPDMM, allowing one to not only cover the immediate operative period, and allow extended coverage until full healing has occurred.

We present studies of candidate drugs for therapeutic intervention in RUNX^+/-^. Yield of megakaryocytes and their agonist responsiveness are the most-straightforward studies to test a drug’s efficacy; however, the xenotransfusion system using shRX-megakaryocytes from healthy adult donor CD34^+^-HSPCs infused into NSG/VWF^R1326H^ mice allows a more detailed look at important aspects of correcting platelet biology, and correcting the hemostatic defects in these mice. A thrombosis challenge as represented by our photochemical injury studies would be one potential preclinical study to correct the bleeding manifestation in patients with FPDMM. Our studies clearly showed a positive effect of RepSox on each defined defect in the megakaryocyte:platelet axis due to RUNX1 deficiency and a strong correction of overall hemostasis. RepSox is thought to inhibit the TGF receptor 1β uptake of TGFβ1^27^, a pathway that we had previously shown to be defective in RUNX1 deficiency; however, RepSox has undergone limited published in vitro and animal studies^27,52^, so it is clearly possible that RepSox targets an alternative pathway. Ongoing studies to define candidate pathways in RUNX1-deficient megakaryocytes following RepSox treatment are underway.

The presented xeno-infusion model takes advantage of prior observations made by our group^24,53^ and others^26^ that marrow-derived megakaryocytes significantly contribute to physiologic platelet production after becoming entrapped in the lungs. It can be applied to the study of other acquired and inherited megakaryocyte/platelet disorders and may provide important in vivo insights into those disorders as well. There are also important limitations to this xeno-infusion model of shRX-megakaryocytes, including the fact that this is a short-term model. Drug exposure in this model is of committed differentiating human hematopoietic cells. The effects of chronic drug exposure on stem and progenitor cells, and the effects of drugs on supporting stromal cells is unexamined. We are presently trying to develop a human xeno-transplantation system beginning with shRX-HSPCs to overcome these two challenges. An additional advantage of developing such a xenotransplant model would be that this system may allow studies of therapeutic interventions in impeding clonal progression to MDS or leukemia.

In summary, we developed a model system using shRX suppression of RUNX1 expression in differentiating CD34^+^-HSPCs to study the qualitative and quantitative defects in the final differentiated megakaryocytes, showing previously unrecognized defects in thrombopoiesis and circulating platelet half-life as well as significant defects in agonist responsiveness. Upon testing the effects of potential therapeutics on differentiating shRX-megakaryocytes, only treatment with RepSox, a TGFβ1 pathway inhibitor, has a clear effect on all the defects in the shRX-megakaryocytes and platelets, and able to correct a hemostatic defect in recipient mice. We propose that this xenotransfusion model may be useful preclinically for drug testing before committing to a clinical trial for correcting the bleeding manifestations of FPDMM.

## Acknowledgments

This work was supported by grants to MP from the RUNX1 Research Program in association with the Alex’s Lemonade Foundation and from the NHLBI (R35 HL150698). We thank Nancy A. Speck from the University of Pennsylvania for helpful discussions and Dr. Douglas B. Cines from the University of Pennsylvania for reviewing the manuscript.

## Author contributions

KL carried out most of the described studies with advice and assistance in both the in vitro and murine xenotransfusion studies from HA. KL also designed new experimental design and prepared the first draft of the paper. BE developed the shRNA study and carried out initial drug therapy studies. MP developed the initial overall study and did data interpretation with KL as well as assist KL in preparation of the manuscript.

## References

1. Okuda T, van Deursen J, Hiebert SW, Grosveld G, Downing JR. AML1, the target of multiple chromosomal translocations in human leukemia, is essential for normal fetal liver hematopoiesis. Cell. 1996;84(2):321–330.

2. Wang Q, Stacy T, Miller JD, et al. The CBFbeta subunit is essential for CBFalpha2 (AML1) function in vivo. Cell. 1996;87(4):697–708.

3. Kalev-Zylinska ML, Horsfield JA, Flores MV, et al. Runx1 is required for zebrafish blood and vessel development and expression of a human RUNX1-CBF2T1 transgene advances a model for studies of leukemogenesis. Development. 2002;129(8):2015–2030.

4. Lee TI, Young RA. Transcriptional regulation and its misregulation in disease. Cell. 2013;152(6):1237–1251.

5. Ichikawa M, Asai T, Saito T, et al. AML-1 is required for megakaryocytic maturation and lymphocytic differentiation, but not for maintenance of hematopoietic stem cells in adult hematopoiesis. Nat Med. 2004;10(3):299–304.

6. Sun W, Downing JR. Haploinsufficiency of AML1 results in a decrease in the number of LTR-HSCs while simultaneously inducing an increase in more mature progenitors. Blood. 2004;104(12):3565–3572.

7. Goldfarb AN. Transcriptional control of megakaryocyte development. Oncogene. 2007;26(47):6795–6802.

8. Tijssen MR, Ghevaert C. Transcription factors in late megakaryopoiesis and related platelet disorders. J Thromb Haemost. 2013;11(4):593–604.

9. Yoshida H, Lareau CA, Ramirez RN, et al. The cis-Regulatory Atlas of the Mouse Immune System. Cell. 2019;176(4):897-912.e20.

10. Schlegelberger B, Heller PG. RUNX1 deficiency (familial platelet disorder with predisposition to myeloid leukemia, FPDMM). Semin Hematol. 2017;54(2):75–80.

11. Song WJ, Sullivan MG, Legare RD, et al. Haploinsufficiency of CBFA2 causes familial thrombocytopenia with propensity to develop acute myelogenous leukaemia. Nat Genet. 1999;23(2):166–175.

12. Preudhomme C, Renneville A, Bourdon V, et al. High frequency of RUNX1 biallelic alteration in acute myeloid leukemia secondary to familial platelet disorder. Blood. 2009;113(22):5583–5587.

13. Jongmans MC, Kuiper RP, Carmichael CL, et al. Novel RUNX1 mutations in familial platelet disorder with enhanced risk for acute myeloid leukemia: clues for improved identification of the FPD/AML syndrome. Leukemia. 2010;24(1):242–246.

14. Babushok DV, Bessler M, Olson TS. Genetic predisposition to myelodysplastic syndrome and acute myeloid leukemia in children and young adults. Leuk Lymphoma. 2016;57(3):520–536.

15. Kanagal-Shamanna R, Loghavi S, DiNardo CD, et al. Bone marrow pathologic abnormalities in familial platelet disorder with propensity for myeloid malignancy and germline RUNX1 mutation. Haematologica. 2017;102(10):1661–1670.

16. Cavalcante de Andrade Silva M, Krepischi ACV, Kulikowski LD, et al. Deletion of RUNX1 exons 1 and 2 associated with familial platelet disorder with propensity to acute myeloid leukemia. Cancer Genet. 2018;222-223:32–37.

17. Cai X, Gaudet JJ, Mangan JK, et al. Runx1 loss minimally impacts long-term hematopoietic stem cells. PLoS One. 2011;6(12):e28430.

18. Vo KK, Jarocha DJ, Lyde RB, et al. FLI1 level during megakaryopoiesis affects thrombopoiesis and platelet biology. Blood. 2017;129(26):3486–3494.

19. Connelly JP, Kwon EM, Gao Y, et al. Targeted correction of RUNX1 mutation in FPD patient-specific induced pluripotent stem cells rescues megakaryopoietic defects. Blood. 2014;124(12):1926–1930.

20. Antony-Debré I, Manchev VT, Balayn N, et al. Level of RUNX1 activity is critical for leukemic predisposition but not for thrombocytopenia. Blood. 2015;125(6):930–940.

21. Li Y, Jin C, Bai H, et al. Human NOTCH4 is a key target of RUNX1 in megakaryocytic differentiation. Blood. 2018;131(2):191–201.

22. Estevez B, Borst S, Jarocha D, et al. RUNX-1 haploinsufficiency causes a marked deficiency of megakaryocyte-biased hematopoietic progenitor cells. Blood. 2021;137(19):2662–2675.

23. Ditadi A, Sturgeon CM, Keller G. A view of human haematopoietic development from the Petri dish. Nat Rev Mol Cell Biol. 2017;18(1):56–67.

24. Wang Y, Hayes V, Jarocha D, et al. Comparative analysis of human ex vivo-generated platelets vs megakaryocyte-generated platelets in mice: a cautionary tale. Blood. 2015;125(23):3627–3636.

25. Fuentes R, Wang Y, Hirsch J, et al. Infusion of mature megakaryocytes into mice yields functional platelets. J Clin Invest. 2010;120(11):3917–3922.

26. Lefrançais E, Ortiz-Muñoz G, Caudrillier A, et al. The lung is a site of platelet biogenesis and a reservoir for haematopoietic progenitors. Nature. 2017;544(7648):105–109.

27. Chen J, Tan K, Zhou H, et al. Modifying murine von Willebrand factor A1 domain for in vivo assessment of human platelet therapies. Nat Biotechnol. 2008;26(1):114–119.

28. Adair BD, Alonso JL, van Agthoven J, et al. Structure-guided design of pure orthosteric inhibitors of αIIbβ3 that prevent thrombosis but preserve hemostasis. Nat Commun. 2020;11(1):398.

29. Huang N, Lou M, Liu H, Avila C, Ma Y. Identification of a potent small molecule capable of regulating polyploidization, megakaryocyte maturation, and platelet production. J Hematol Oncol. 2016;9(1):136.

30. Jarocha D, Vo KK, Lyde RB, Hayes V, Camire RM, Poncz M. Enhancing functional platelet release in vivo from in vitro-grown megakaryocytes using small molecule inhibitors. Blood Adv. 2018;2(6):597–606.

31. Furman MI, Liu L, Benoit SE, Becker RC, Barnard MR, Michelson AD. The cleaved peptide of the thrombin receptor is a strong platelet agonist. Proc Natl Acad Sci U S A. 1998;95(6):3082–3087.

32. Rodriguez BAT, Bhan A, Beswick A, et al. A Platelet Function Modulator of Thrombin Activation Is Causally Linked to Cardiovascular Disease and Affects PAR4 Receptor Signaling. Am J Hum Genet. 2020;107(2):211–221.

33. Francischetti IM, Ghazaleh FA, Reis RA, Carlini CR, Guimarães JA. Convulxin induces platelet activation by a tyrosine-kinase-dependent pathway and stimulates tyrosine phosphorylation of platelet proteins, including PLC gamma 2, independently of integrin alpha IIb beta 3. Arch Biochem Biophys. 1998;353(2):239–250.

34. Kanaji S, Kanaji T, Furihata K, Kato K, Ware JL, Kunicki TJ. Convulxin binds to native, human glycoprotein Ib alpha. J Biol Chem. 2003;278(41):39452–39460.

35. Kinlough-Rathbone RL, Rand ML, Packham MA. Rabbit and rat platelets do not respond to thrombin receptor peptides that activate human platelets. Blood. 1993;82(1):103–106.

36. Huang H, Woo AJ, Waldon Z, et al. A Src family kinase-Shp2 axis controls RUNX1 activity in megakaryocyte and T-lymphocyte differentiation. Genes Dev. 2012;26(14):1587–1601.

37. Angers-Loustau A, Hering R, Werbowetski TE, Kaplan DR, Del Maestro RF. SRC regulates actin dynamics and invasion of malignant glial cells in three dimensions. Mol Cancer Res. 2004;2(11):595–605.

38. Angell RM, Atkinson FL, Brown MJ, et al. N-(3-Cyano-4,5,6,7-tetrahydro-1-benzothien-2-yl)amides as potent, selective, inhibitors of JNK2 and JNK3. Bioorg Med Chem Lett. 2007;17(5):1296–1301.

39. Olsauskas-Kuprys R, Zlobin A, Osipo C. Gamma secretase inhibitors of Notch signaling. Onco Targets Ther. 2013;6:943–955.

40. Glembotsky AC, Bluteau D, Espasandin YR, et al. Mechanisms underlying platelet function defect in a pedigree with familial platelet disorder with a predisposition to acute myelogenous leukemia: potential role for candidate RUNX1 targets. J Thromb Haemost. 2014;12(5):761–772.

41. Rao AK, Poncz M. Defective acid hydrolase secretion in RUNX1 haplodeficiency: Evidence for a global platelet secretory defect. Haemophilia. 2017;23(5):784–792.

42. Secchiero P, Voltan R, Rimondi E, et al. The γ-secretase inhibitors enhance the anti-leukemic activity of ibrutinib in B-CLL cells. Oncotarget. 2017;8(35):59235–59245.

43. Cheng M, Lv X, Zhang C, et al. DNMT1, a Novel Regulator Mediating mTORC1/mTORC2 Pathway-Induced NGF Expression in Schwann Cells. Neurochem Res. 2018;43(11):2141–2154.

44. Santini V, Valcárcel D, Platzbecker U, et al. Phase II Study of the ALK5 Inhibitor Galunisertib in Very Low-, Low-, and Intermediate-Risk Myelodysplastic Syndromes. Clin Cancer Res. 2019;25(23):6976–6985.

45. Xiao X, Lai W, Xie H, et al. Targeting JNK pathway promotes human hematopoietic stem cell expansion. Cell Discov. 2019;5:2.

46. Varricchio L, Iancu-Rubin C, Upadhyaya B, et al. TGF-β1 protein trap AVID200 beneficially affects hematopoiesis and bone marrow fibrosis in myelofibrosis. JCI Insight. 2021;6(18):e145651.

47. Boitano AE, de Lichtervelde L, Snead JL, Cooke MP, Schultz PG. An image-based screen identifies a small molecule regulator of megakaryopoiesis. Proc Natl Acad Sci U S A. 2012;109(35):14019–14023.

48. Javarappa KK, Tsallos D, Heckman CA. A Multiplexed Screening Assay to Evaluate Chemotherapy-Induced Myelosuppression Using Healthy Peripheral Blood and Bone Marrow. SLAS Discov. 2018;23(7):687–696.

49. Gaur M, Kamata T, Wang S, Moran B, Shattil SJ, Leavitt AD. Megakaryocytes derived from human embryonic stem cells: a genetically tractable system to study megakaryocytopoiesis and integrin function. J Thromb Haemost. 2006;4(2):436–442.

50. Brown AL, Arts P, Carmichael CL, et al. RUNX1-mutated families show phenotype heterogeneity and a somatic mutation profile unique to germline predisposed AML. Blood Adv. 2020;4(6):1131–1144.

51. Churpek JE, Pyrtel K, Kanchi KL, et al. Genomic analysis of germ line and somatic variants in familial myelodysplasia/acute myeloid leukemia. Blood. 2015;126(22):2484–2490.

52. Ide M, Jinnin M, Tomizawa Y, et al. Transforming growth factor β-inhibitor Repsox downregulates collagen expression of scleroderma dermal fibroblasts and prevents bleomycin-induced mice skin fibrosis. Exp Dermatol. 2017;26(11):1139–1143.

53. Fuentes R, Wang Y, Hirsch J, et al. Infusion of mature megakaryocytes into mice yields functional platelets. J Clin Invest. 2010;120(11):3917–3922. doi:10.1172/JCI43326

