## supplemental figures for "RUNX1-deficient human megakaryocytes demonstrate thrombopoietic and platelet half-life and functional defects: Therapeutic implications"

#### Supplemental material and method

*Platelet isolation*

Five ml of human peripheral blood was drawn into a 15 ml centrifuge tube containing 500 μl of 3.8% sodium citrate using a 21 G syringe needle from healthy adult donors. Only supernatant (platelet-rich plasma, PRP) was used after centrifugation of the whole blood at 350xg with acceleration set at 3 and deceleration at 1 done for 10 minutes at room temperature. Platelets were enumerated using an automatic cell counter (HEMAVET, Drew Scientific). PRP was centrifuged at 2,000xg for 10 minutes at room temperature to pellet platelets.

*Sorting for shRNA-expressing CD34$^+$-HSPCs*

At 48 hours post-lentiviral transduction into CD34$^+$-HSPCs, cells were centrifuged and resuspended in cold sorting buffer (1% FBS and 2 mM EDTA in PBS) at 1X10$^7$ cells/ml on ice. Only mCherry$^+$ populations were isolated by FACS Jazz (Becton-Dickinson) with a 100μ nozzle as described under RNA isolation and RT-PCR[1]. After sorting the cells were replated in differentiation medium.

*RNA isolation and RT-qPCR*

At 72 hours after transduction of lentiviral shRNA, shNT or shRX-CD34$^+$ cells were sorted for mCherry$^+$ populations as described in Sorting for shRNA-expressing CD34$^+$-HSPCs. Total RNAs were isolated with Trizol (Invitrogen) from 1X10$^5$ sorted cells. cDNAs were generated by reverse transcription with iScript (Bio-Rad, 170-8840) including oligo (dT) primers and human *RUNX1* (*RUNX1* forward primer: 5'-AGTGGAAGAGGGAAAAGC-3', reverse primer: 5'-ATCCACTGTGATTTTGATGG-3') mRNA levels were quantified by using real-time PCR with SYBR green dye on an ABI Prism 7000 system. For an internal control, human *GAPDH* primers were used (*GAPDH* forward primer: 5'-TGGTATCGTGGAAGGACTCATGAC-3', reverse primer: 5'-ATGCCAGTGAGCTTCCCGTTCAGC-3'). Total mRNA levels of human *RUNX1* were normalized for *GAPDH* levels.

*Ploidy study*

mCherry$^+$- megakaryocytes were fixed in 70% ethanol on ice for 15 min. After washing twice in BD Perm/Wash buffer (Becton-Dickinson, 51-2091KZ), cells were stained in 1 μg/ml propidium iodide (Millipore, 537059) for 15 minutes on ice in dark. Cells were washed twice in cold PBS and then analyzed by a CytoFLEX 6 (Becton-Dickinson).

*Intracellular and surface immunostaining for flow*

Either shNT- or shRX-megakaryocytes (Day 11) were fixed in 4.2% formaldehyde in PBS (BD Cytofix/Cytoperm buffer, 51-2091KZ) on ice for 15 minutes and then washed them twice in BD Perm/Wash buffer. Fixed cells were incubated with anti-RUNX1 antibody (1:100 dilution, Santa Cruz SC-365644) for 30 minutes on ice and then washed them twice. A fluorescent second antibody, FITC-conjugated goat anti-mouse IgG (1:100 dilution, Becton-Dickinson 555988), was incubated on ice for 30 minutes and then washed them twice. For a background control, only the second antibody was incubated. After washing twice, cells were analyzed by a CytoFLEX 6 (Becton-Dickinson).

To measure human erythroid and myeloid populations in $CD34^+$-megakaryocyte differentiation in $RUNX1^{Lo}$ cells were stained with APC-labeled polyclonal rabbit anti-CD235a for human erythroid and APC/Cy7-labeled polyclonal rabbit anti-CD11b for human myeloid.

*Counting absolute number of $CD34^+$-megakaryocytes*

For the drug screening and studies of Mk yield, we started culturing with $10^6$ cells of $CD34^+$ and sorted only $mCherry^+$ cells (~30%) at Day 4. The same number of sorted cells ($10^4$ per well of 96-well plate) were then plated in 150 μl of differentiation medium per well. On Day 11 or Day 13-14, the absolute number of mature megakaryocytes were calculated by using CountBright beads (Invitrogen, C36950). The absolute number of $CD34^+$-Mk was calculated by the followed equitation: absolute count (cells/μl) = cell count/counting beads count x counting beads concentration (beads/μl).

*Drugs*

RepSox (RS, Stemcell, 72392), Galunisertib (Selleckchem, S2230), DAPT (MedChemExpress, HY-13027), Avagacestat (Selleckchem, BMS-708163), JIN8 (Selleckchem, S4901), JIN10 (Selleckchem, S7508), MHY (Selleckchem, 326914-06-1), PP2 (Selleckchem, S7008), Imatinib (MedChemExpress, HY-50946), Nilotinib (MedChemExpress, HY-10159), Ruxolitinib (Selleckchem, S1378)

#### Supplemental figure legends

**Figure S1. The lentiviral shRNA constructs and sorting shRNA-expressing cells.**

(**A**) Lentiviral-shRNA construct with a shNT or shRX sequence inserted are depicted in constructs downstream of a murine stem-cell virus (MSCV) promoter in a microRNA (mir) scaffold. mCherry gene is expressed as a marker of transduction[1]. Other essential elements of pCL20.MSCV.shLuc.PGK.mCherry vectors are shown. LTR = long terminal repeat. (**B**) At 48 hours after lentiviral transduction, only mCherry$^+$ cells were sorted with the indicated gating (red boxes). After sorting, the collected populations were confirmed (bottom panels) by flow. Both shNT and shRX samples were >90% mCherry$^+$ cells post-sorting.

**Figure S2. Representative flow cytometry showing high level of continued mCherry positive cells during differentiation from Day 4 infection at least until Day 11.**

Flow cytometric studies of mCherry positivity of untransfected, and shNT- and shRX-transfected hematopoietic cells on Day 4 (top) and Day 11(bottom).

**Figure S3. Relative level of RUNX1 mRNA and protein yield in RUNX1 knockdown CD34$^+$-megakaryocytes.**

(**A**) mRNA levels in CD34$^+$ cells at 96 hours after infection with lentiviral shNT or shRX measured by RT-qPCR were determined. Those levels were normalized to human GAPDH mRNA levels as an internal control and then expressed as relative to the shNT-megakaryocyte level. Grey dashed line is the 100% relative of the shNT-megakaryocytes. N = 3 per arm. Mean ± 1 SD is shown. P-value is calculated by using a two-tailed t-test. (**B**) Same as (A), but measurement of RUNX1 protein level by flow cytometry with an additional isotype control arm. N=3 per arm. P-value is calculated by using a two-tailed t-test. (C) Mean ± 1 SD for absolute number of mature CD41$^+$CD42$^+$ megakaryocyte yield from $10^6$ mCherry$^+$ progenitors after shNT or shRX lentiviral transduction and beginning with $10^4$ mCherry$^+$ progenitor cells per well. Studies of shRX-megakaryocytes were also done in the presence of 1 μM RS as well. N = 3 per experiment. P-values were calculated using two-way Student t test.

**Figure S4. Effects of RUNX1 knockdown on progenitors and definitive lineages and on megakaryocyte ploidy.**

(**A**) Representative flow cytometric data on the effects of RUNX1 knockdown and subsequent exposure top 1 μM RS on CD41 and CD42 expression on Day 11 of mCherry+ cells. (**B**) Same as in (A) but showing mean ± 1 SD for immature and mature megakaryocytes and also for human erythroid (CD235) and myeloid (CD11b) markers. N = 3 per arm. P-value is calculated by using a two-tailed t-test. * = p<0.05 and ** = p<0.01 comparing yield of indicated cell population after shRX exposure to shNT control. (**C**) Representative flow cytometric data in Day 4 (left) and 11 (middle and right) after propidium iodide staining for ploidy. Cells on right had been exposed to RepSox as described.

**Figure S5. Agonist-responsiveness test in Day 14 RUNX1$^{Lo}$ CD34$^{+}$-megakaryocytes.**

(**A**) Schematic of agonist-induced megakaryocytes (and also platelet (Plts)) activation. Different kinds of agonists can activate megakaryocytes or platelets through their cognate membrane receptors as indicated. Studies focused on thrombin or the human platelet-specific protease-activating receptor 1 peptide TRAP[2], or convulxin as a collagen agonist[3, 4] for glycoprotein VI. As a result of activation of megakaryocytes or platelets, the cells degranulate and express surface P-selectin[5, 6]. (**B**) On Day 11 of differentiation, shNT- or shRX-megakaryocytes were exposed to various doses of thrombin as indicated by darker colors and then the level of surface expressed P-selectin was measured. mCherry$^{-}$ samples are in gray, mCherry$^{+}$ samples are in blue (shNT) or red (shRX). As controls, undifferentiated CD34-HSPCs (Undiff) or uninfected populations (mCherry$^{-}$ population) were examined (gray bars). Mean ± 1 SD shown. N = 3. Mean ± 1 SD are shown. * = P ≤ 0.05, ** = P ≤ 0.01, and *** = P ≤ 0.001, and **** = P ≤ 0.0001. P- values were calculated by one-way ANOVA comparing shRX to shNT comparable doses. (**C**) and (**D**) are as shown in (B), but after exposure to varied doses of TRAP (C) or convulxin (D).

**Figure S6. Initial blood flow in the hemostatic injury model**

For the control of *in vivo* thrombus formation studies as in Figure 7, initial blood flow was measured for each group. Mean ± 1 SD shown. N = 4-6 per arm. ns = not significant. P-values were calculated by one-way ANOVA comparing indicated pairs.

**Figure S7. Drug screening to rescue RUNX1-deficient Day 11 Mks.**

Same as Figure 4, but for Day 11 Mks with (**A**) being studies with shNT-megakaryocytes and (**B**) with shRX-megakaryocytes. Mean ± 1 SD shown. The dashed line represents yield of megakaryocytes from differentiated from uninfected HSPCs not exposed to any drug. N = 3 per arm. * = P ≤ 0.05, *** = P ≤ 0.001, and **** = P ≤ 0.0001. P-values were calculated by one-way ANOVA compared to each DMSO control.

**References**


1. Estevez B, Borst S, Jarocha D, et al. RUNX-1 haploinsufficiency causes a marked deficiency of megakaryocyte-biased hematopoietic progenitor cells. *Blood*. 2021;137(19):2662-2675.
2. Kinlough-Rathbone RL, Rand ML, Packham MA. Rabbit and rat platelets do not respond to thrombin receptor peptides that activate human platelets. *Blood*. 1993;82(1):103-106.
3. Francischetti IM, Ghazaleh FA, Reis RA, Carlini CR, Guimarães JA. Convulxin induces platelet activation by a tyrosine-kinase-dependent pathway and stimulates tyrosine phosphorylation of platelet proteins, including PLC gamma 2, independently of integrin alpha IIb beta 3. *Arch Biochem Biophys*. 1998;353(2):239-250.
4. Kanaji S, Kanaji T, Furihata K, Kato K, Ware JL, Kunicki TJ. Convulxin binds to native, human glycoprotein Ib alpha. *J Biol Chem*. 2003;278(41):39452-39460.
5. Hattori R, Hamilton KK, Fugate RD, McEver RP, Sims PJ. Stimulated secretion of endothelial von Willebrand factor is accompanied by rapid redistribution to the cell surface of the intracellular granule membrane protein GMP-140. J Biol Chem. 1989;264(14):7768-7771.
6. Disdier M, Morrissey JH, Fugate RD, Bainton DF, McEver RP. Cytoplasmic domain of P-selectin (CD62) contains the signal for sorting into the regulated secretory pathway. Mol Biol Cell. 1992;3(3):309-321.

### Figure S1

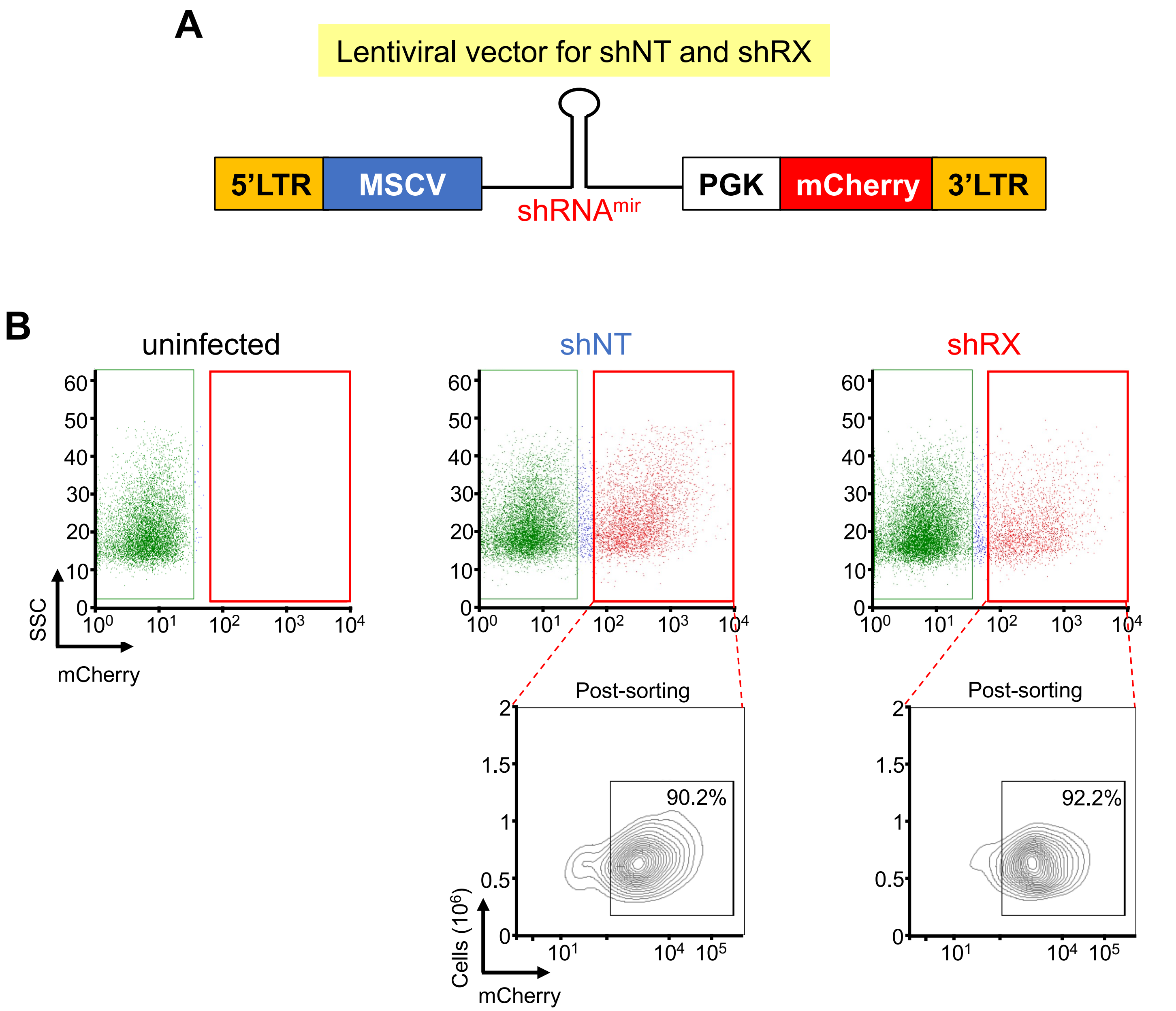
A
Lentiviral vector for shNT and shRX
5'LTR
MSCV
shRNAmir
PGK
mCherry
3'LTR
B
uninfected
shNT
shRX
SSC
mCherry
Post-sorting
90.2%
92.2%
Cells (10^6)
mCherry

### Figure S2

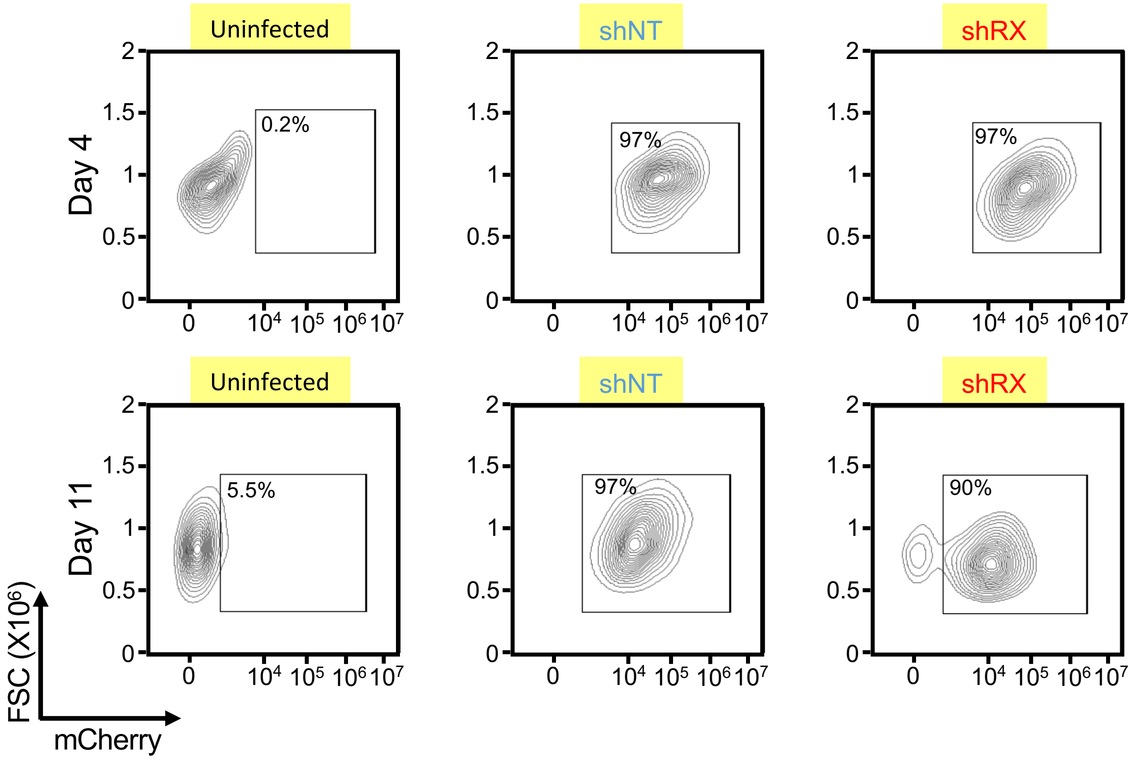

Uninfected
shNT
shRX
Day 4
Day 11
0.2%
97%
97%
5.5%
97%
90%
2
1.5
1
0.5
0
0
10⁴ 10⁵ 10⁶ 10⁷
FSC (X10⁶)
mCherry

### Figure S3

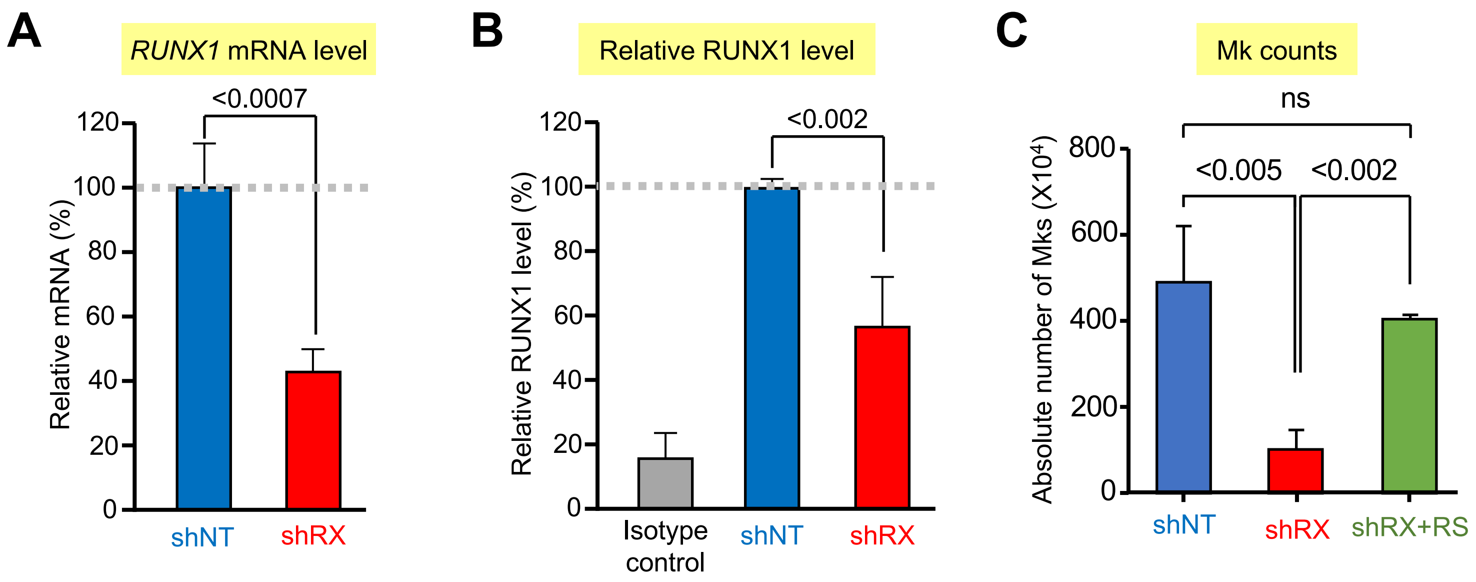
A
RUNX1 mRNA level
<0.0007
Relative mRNA (%)
120
100
80
60
40
20
0
shNT
shRX
B
Relative RUNX1 level
<0.002
Relative RUNX1 level (%)
120
100
80
60
40
20
0
Isotype control
shNT
shRX
C
Mk counts
ns
<0.005
<0.002
Absolute number of Mks (X10⁴)
800
600
400
200
0
shNT
shRX
shRX+RS

### Figure S4

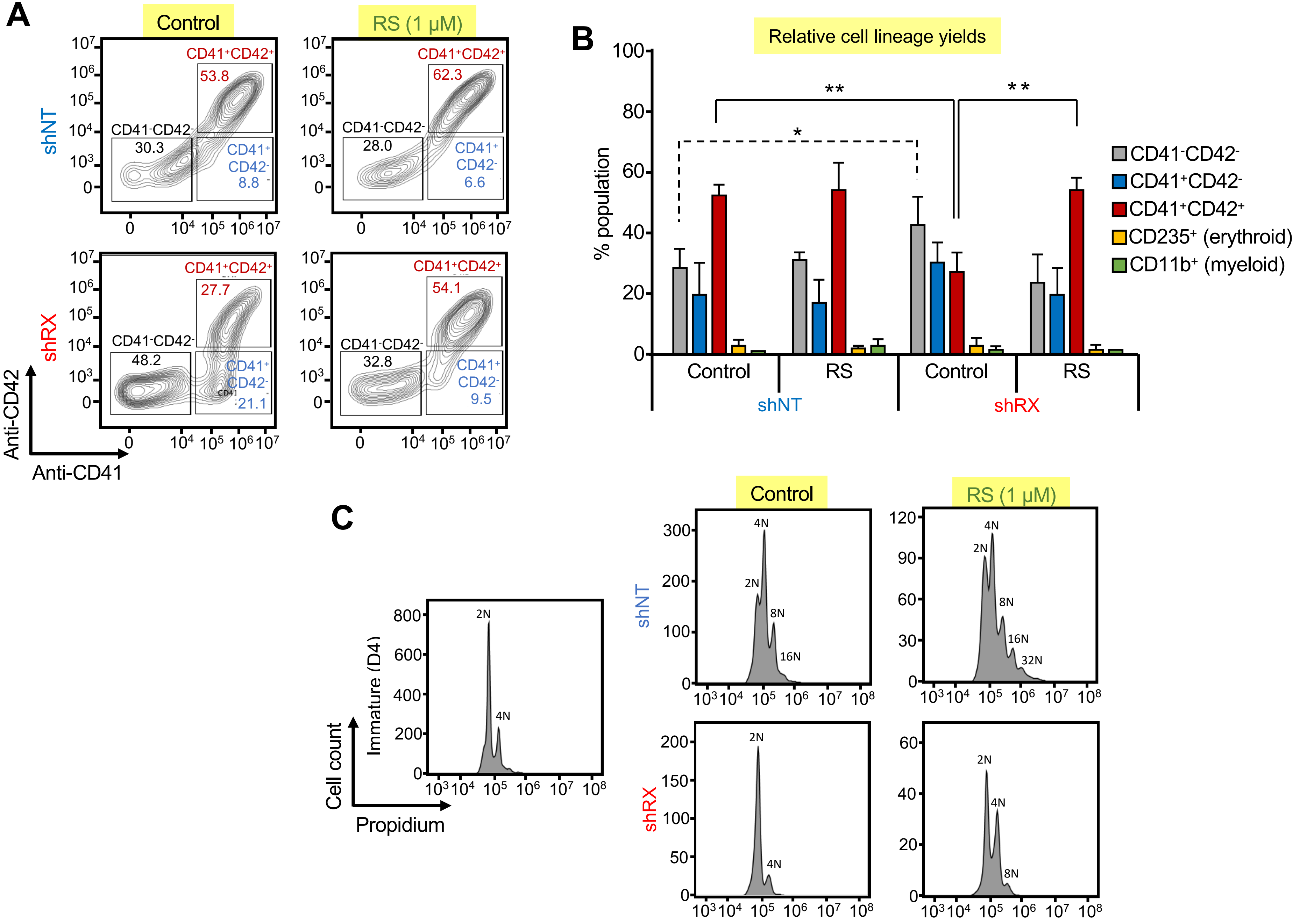
A
Control
RS (1 μM)
shNT
shRX
CD41+CD42+
53.8
62.3
27.7
54.1
CD41-CD42-
30.3
28.0
48.2
32.8
CD41+ CD42-
8.8
6.6
21.1
9.5
Anti-CD42
Anti-CD41
B
Relative cell lineage yields
% population
Control
RS
shNT
shRX
CD41-CD42-
CD41+CD42-
CD41+CD42+
CD235+ (erythroid)
CD11b+ (myeloid)
C
Cell count
Immature (D4)
Propidium
2N
4N
8N
16N
32N

### Figure S5

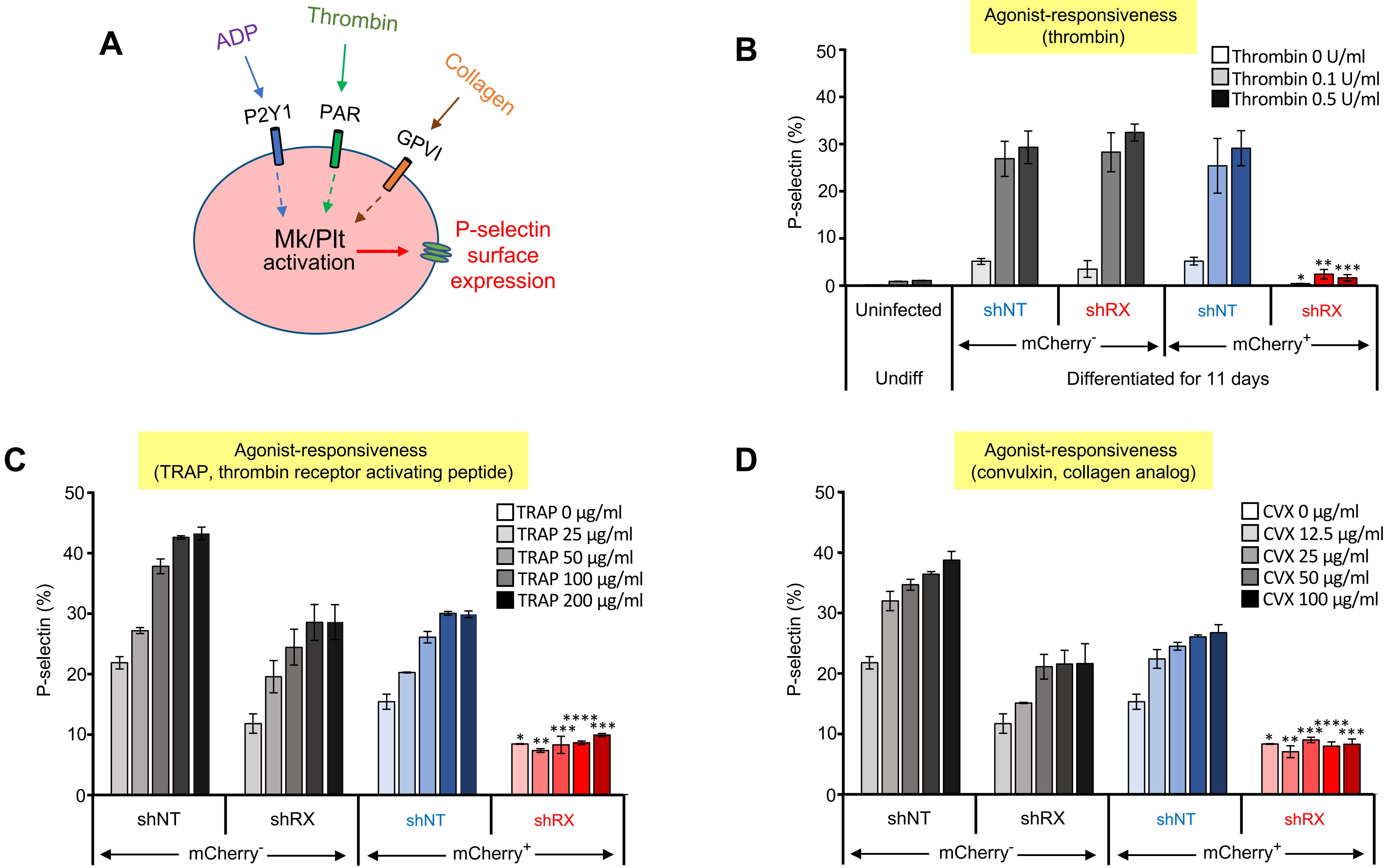
A
ADP
Thrombin
Collagen
P2Y1
PAR
GPVI
Mk/Plt activation
P-selectin surface expression
B
Agonist-responsiveness (thrombin)
Thrombin 0 U/ml
Thrombin 0.1 U/ml
Thrombin 0.5 U/ml
P-selectin (%)
Uninfected
shNT
shRX
shNT
shRX
mCherry-
mCherry+
Undiff
Differentiated for 11 days
C
Agonist-responsiveness (TRAP, thrombin receptor activating peptide)
TRAP 0 μg/ml
TRAP 25 μg/ml
TRAP 50 μg/ml
TRAP 100 μg/ml
TRAP 200 μg/ml
P-selectin (%)
shNT
shRX
shNT
shRX
mCherry-
mCherry+
D
Agonist-responsiveness (convulxin, collagen analog)
CVX 0 μg/ml
CVX 12.5 μg/ml
CVX 25 μg/ml
CVX 50 μg/ml
CVX 100 μg/ml
P-selectin (%)
shNT
shRX
shNT
shRX
mCherry-
mCherry+

### Figure S6

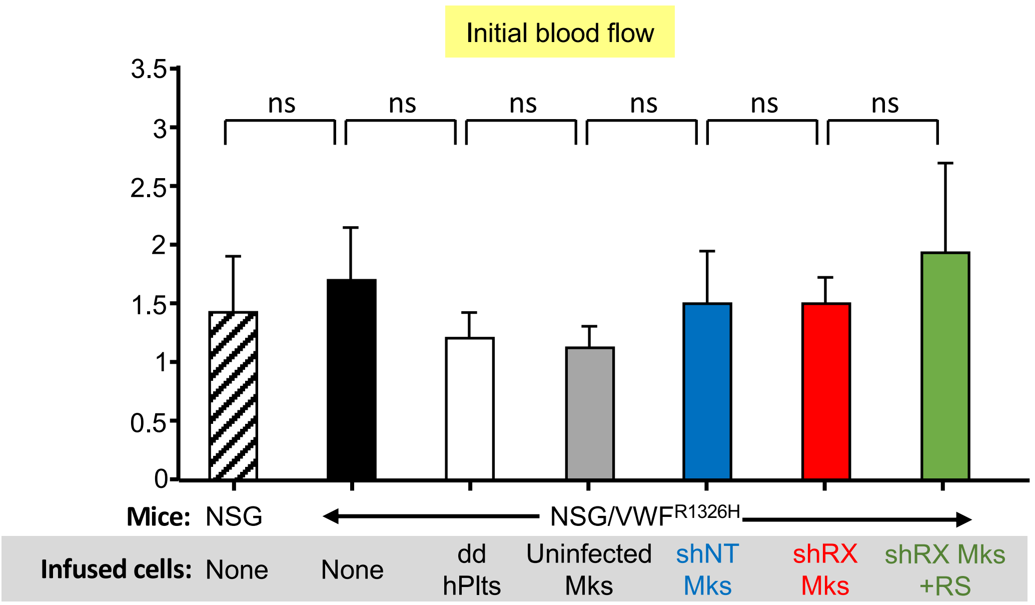
Initial blood flow
3.5
3
2.5
2
1.5
1
0.5
0
ns
ns
ns
ns
ns
ns
Mice: NSG
NSG/VWF R1326H
Infused cells: None
None
dd hPlts
Uninfected Mks
shNT Mks
shRX Mks
shRX Mks +RS

### Figure S7

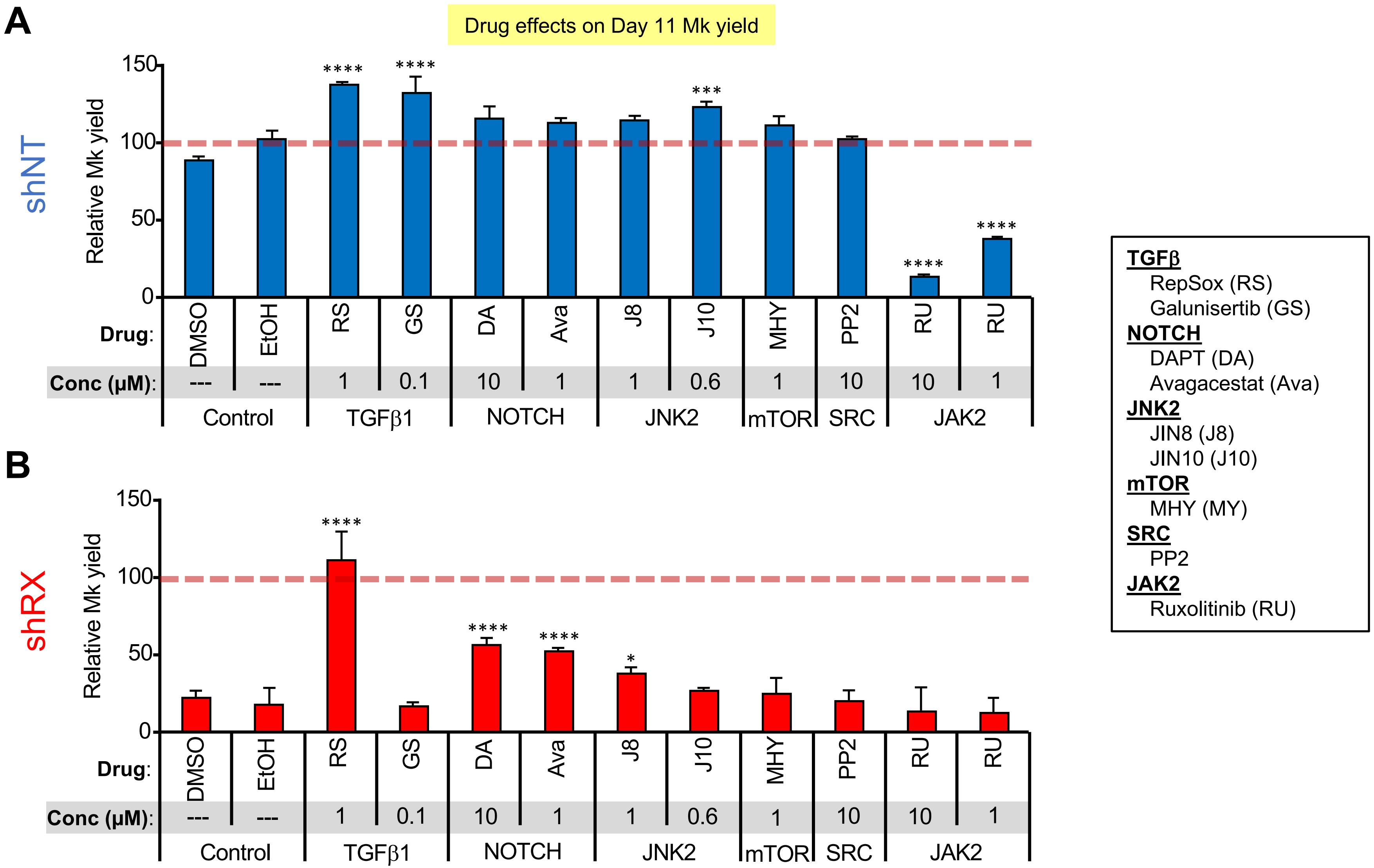
A
Drug effects on Day 11 Mk yield
shNT
Relative Mk yield
150
100
50
0
****
****
***
****
****
Drug:
DMSO
EtOH
RS
GS
DA
Ava
J8
J10
MHY
PP2
RU
RU
Conc (μM):
---
---
1
0.1
10
1
1
0.6
1
10
10
1
Control
TGFβ1
NOTCH
JNK2
mTOR
SRC
JAK2
B
shRX
Relative Mk yield
150
100
50
0
****
****
****
*
Drug:
DMSO
EtOH
RS
GS
DA
Ava
J8
J10
MHY
PP2
RU
RU
Conc (μM):
---
---
1
0.1
10
1
1
0.6
1
10
10
1
Control
TGFβ1
NOTCH
JNK2
mTOR
SRC
JAK2
TGFβ
RepSox (RS)
Galunisertib (GS)
NOTCH
DAPT (DA)
Avagacestat (Ava)
JNK2
JIN8 (J8)
JIN10 (J10)
mTOR
MHY (MY)
SRC
PP2
JAK2
Ruxolitinib (RU)
